# A novel peptide modulator of a two-component system revealed by the specific activation of a small RNA in *Enterobacteriaceae*

**DOI:** 10.64898/2026.03.20.713024

**Authors:** Jade Mathis de Fromont, Anaïs Brosse, Fanny Quenette, Maude Guillier

## Abstract

Small regulatory RNAs (sRNAs) are major post-transcriptional regulators in bacteria and, together with transcriptional regulators such as the two-component systems (TCSs), participate in the rapid adaptation of these microorganisms to changing environments. Several examples of paralogous sRNAs with overlapping functions have been reported, that could in theory integrate different environmental cues. Consistent with this idea, we have identified the acid-responsive RstB-RstA two-component system, important for virulence of multiple bacterial species, as a specific multicopy activator of the *Escherichia coli* OmrB sRNA, but not of the paralogous sRNA OmrA. Further characterization of this regulation unexpectedly revealed the *asr-ydgU* operon, itself a target of RstB-RstA, as a dual modulator of this TCS via two opposite effects. First, the 27 aminoacids YdgU small protein exerts a negative feedback by directly interacting with RstB and, second, Asr in contrast mediates a positive feedback on RstB-RstA activity via a not completely elucidated mechanism. These results provide a new example of retro-control of a TCS, here RstB-RstA, by one of its direct targets. They further highlight the major role of small proteins in controlling TCS activity and *ydgU* was thus renamed *samT*, for Small Acid-responsive Modulator of the RstB-RstA TCS.

## INTRODUCTION

Control of gene expression allows bacterial adaptation to multiple environments and plays a key role in the ability of these microorganisms to successfully colonize various habitats. Multiple regulators acting at different levels are involved in this process. Among these, two-components systems (TCSs) and small RNAs (sRNAs), acting mostly at the transcriptional and post-transcriptional level, respectively, are widespread in bacteria and participate in the rapid adaptive response to various external and internal cues. TCSs typically respond to specific signals via the autophosphorylation of a sensor kinase, usually located at the membrane, which then transfers the phosphate group to a conserved aspartate residue of a cognate response regulator. The resulting phosphorylation of the response regulator most often leads to transcriptional regulation by increasing its affinity for specific DNA motifs (1). In contrast, the majority of sRNA regulators act post-transcriptionally via imperfect base-pairing to multiple mRNA-targets, thereby affecting their translation and/or stability (2).

Importantly, transcriptional and post-transcriptional controls are often integrated in mixed regulatory circuits that combine multiple regulators. One of many examples is provided by the control of sRNAs transcription by TCSs, while on the other hand sRNAs can base-pair to mRNAs encoding TCS proteins to control their expression. This is notably the case for well-studied TCSs such as PhoQ-PhoP or EnvZ-OmpR that activate transcription of the MgrR and OmrA/B sRNAs, respectively (3, 4). In turn, these two TCSs are also targeted by sRNAs that repress their expression: MicA and GcvB sRNAs base-pair to the *phoP* translation initiation region of the *phoPQ* operon (5, 6) while the OmrA/B sRNAs base-pair to *ompR* translation initiation region in *ompR-envZ*. This last control therefore results in a feedback control of the EnvZ-OmpR TCS by the OmrA/B sRNAs (4) (Fig. 1A).

**Figure 1.**
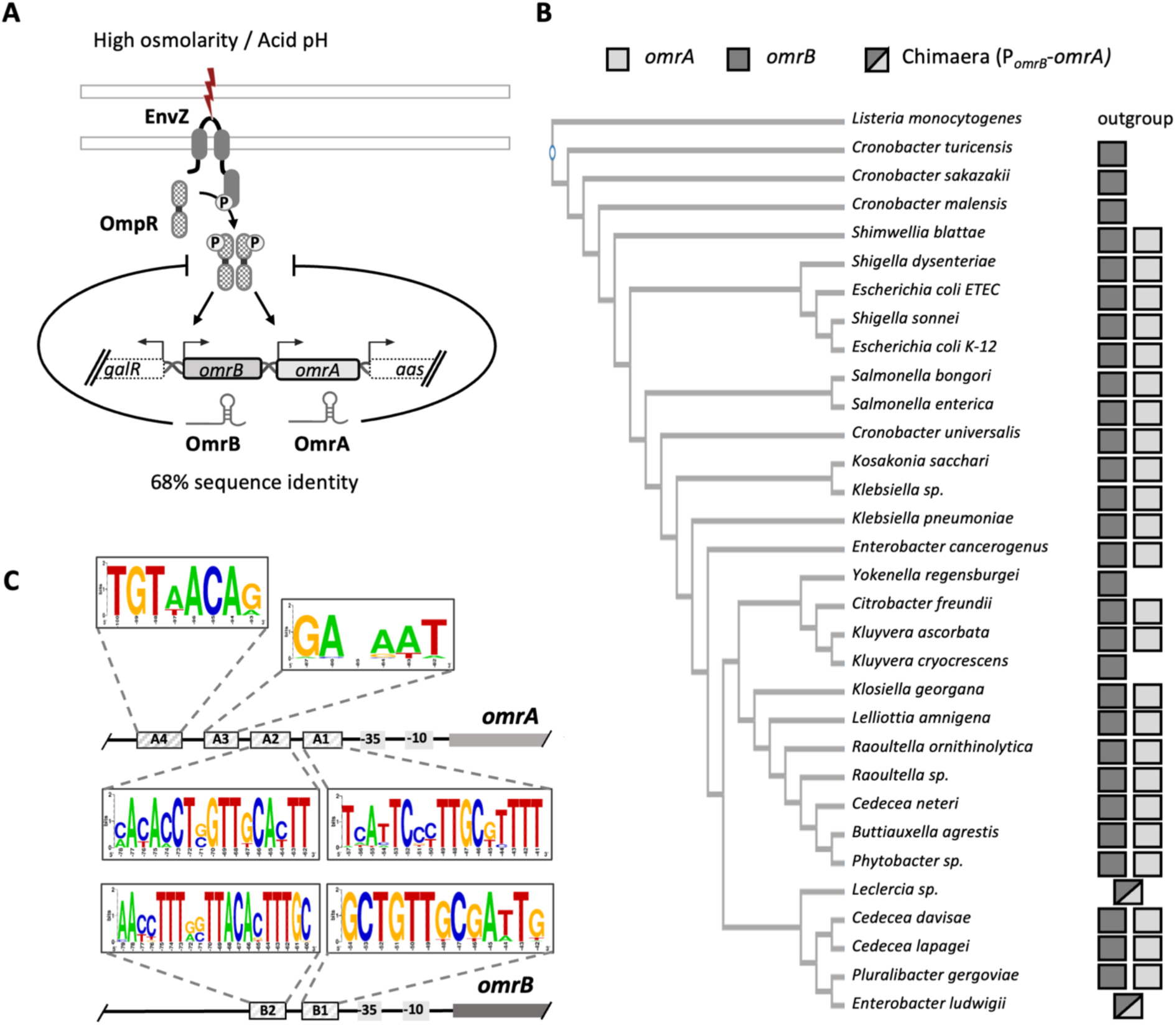
Conservation of the OmrA and OmrB sRNAs in *Enterobacteriaceae*. (A) Scheme of the *E. coli omrAB* genomic organization and known transcriptional control of these sRNAs by the EnvZ-OmpR TCS. The negative post-transcriptional regulation of *ompR* by the OmrA and OmrB sRNAs (13) is schematized by the lines ending with bars. (B) Detection of *omrA* and/or *omrB* sequences in the *aas-galR* intergenic region in the indicated bacteria. The chimeara correspond to a hybrid sequence carrying the *omrB* promoter followed by the *omrA* transcribed sequence. The possibility that another *omr* gene copy could be present on the genome, at a different locus, is not addressed here. (C) Conserved sequences in the *omrA* or *omrB* promoter regions, highlighting the presence of several conserved boxes. The detailed alignment is shown in Fig. S1.

**Figure 2.**
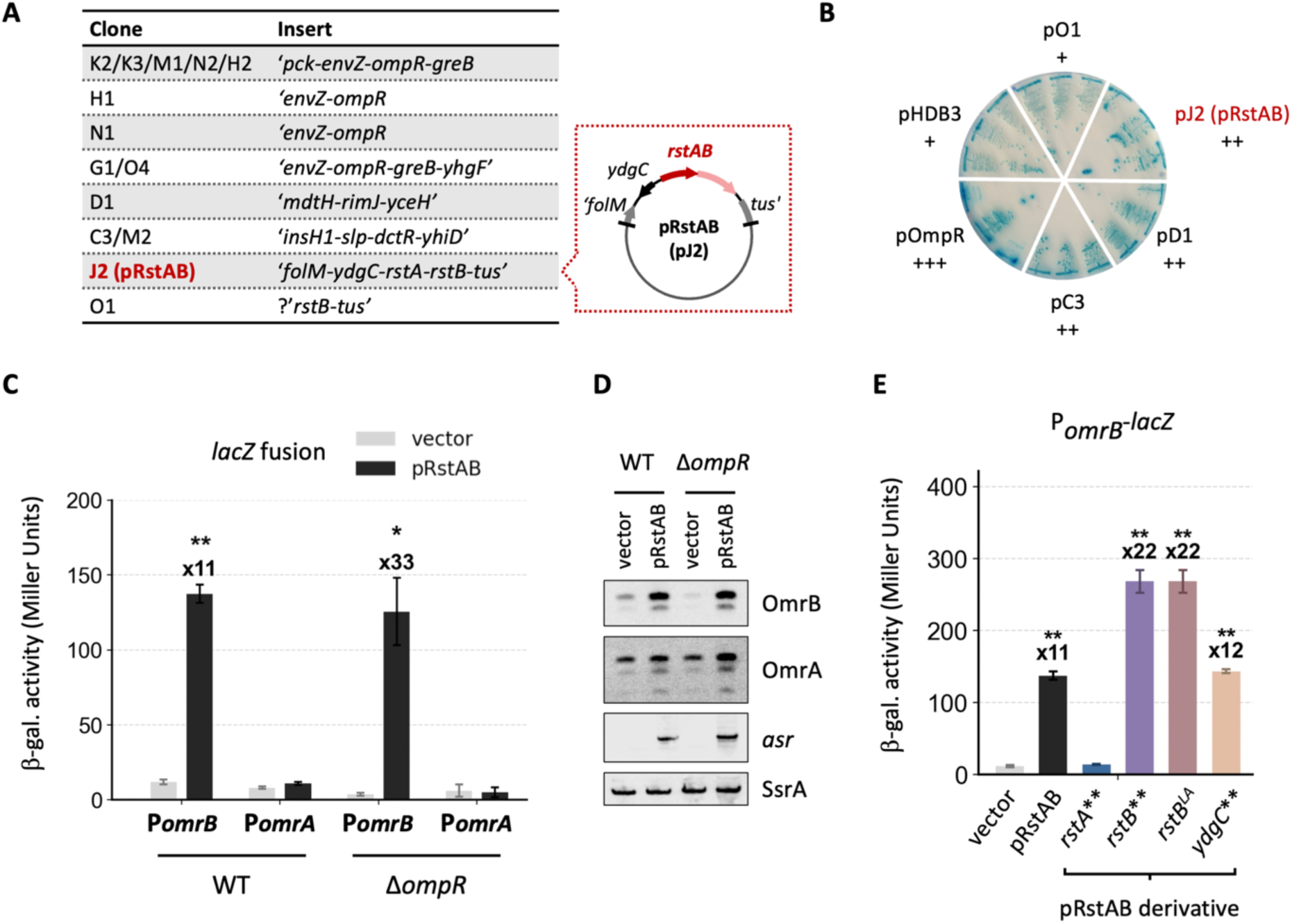
The RstB-RstA two-component system is a specific multicopy activator of *omrB* transcription. (A) Summary of the genomic DNA fragments found on plasmids that increase *omrB* transcription. The inserts from the plasmids shown on the same line are identical. The “?” in plasmid pO1 indicates that the insert boundary could not be mapped, most likely because of plasmid rearrangement. (B) Phenotypes of a strain carrying the P*_omrB_*-*lacZ* fusion and deleted of the *ompR* chromosomal gene (strain MG1812) transformed with several of the plasmids identified as activators in the genetic screen, on a minimal A glucose agar plate, supplemented with 1 mM MgSO4, X-gal 0.002% and ampicillin. pHDB3 is the empty vector and pOmpR the activating plasmid identified previously (12). The “+” signs denote the intensity of the blue color of the colonies when transformed by the indicated plasmid. (C) The β-galactosidase activity of P*_omrA_*- and P*_omrB_*-*lacZ* fusions was measured in the presence of the pRstAB plasmid or the pHDB3 vector control, in an *ompR^+^* or Δ*ompR* background, in minimal A glucose medium, supplemented with ampicillin. Strains used in this experiment are JM2110, JM2111, JM2118 and JM2115. The β-galactosidase activities are the average of three independent experiments and error bars indicate the standard deviation. Statistical significance of the results was assessed by calculation of the p-value through a heteroscedastic two-tails t test (* for p- values ≤0.05, ** for p-values ≤0.01). (D) Northern blot analysis of OmrA and OmrB sRNAs, using total RNA extracted from the cultures of strains JM2110 and JM2111 used in panel (C). The *asr* mRNA was probed on the same membrane as a control for *rstAB* overexpression, and SsrA as a loading control. (E) The effect of several derivatives of the pRstAB plasmid was analyzed on *omrB* transcription using the P*_omrB_*-*lacZ* as a read-out (strain JM2110) as described in (C). These derivatives carry stop codons in *rstA*, *rstB* or *ydgC* early coding sequence (indicated by **), or the *rstB-LA* mutation that prevents phosphotransfer to RstA *in vitro* (37).

In addition to this post-transcriptional control of TCS-encoding genes, the activity of TCSs can also be subject to regulation by so called modulator proteins, some of which are small proteins (less than 50 aminoacids (aa)). Again, examples for such regulation have been reported for EnvZ-OmpR, with the MzrA membrane protein and YoaI small protein (7, 8), or for PhoQ-PhoP, with the SafA, MgrB and UgtS/UgtL proteins (9–11). Hence, multiple pathways participate in the precise control of TCS synthesis and activity, highlighting their importance for bacterial physiology.

The sRNA-mediated feedback control of the EnvZ-OmpR TCS relies on OmrA and OmrB, two paralogous sRNAs that were identified based on their conservation in enterobacteria and their ability to bind the Hfq chaperone. Their 5’ and 3’-end sequences are the most conserved regions and are also almost identical between these two sRNAs, while the central region differs between OmrA and OmrB (12). The conserved 5’-end of both OmrA and OmrB base-pairs to multiple targets, that are thus common for these two sRNAs, such as the *ompR-envZ* operon mentioned above. Other common targets reported so far, all negatively regulated, encode outer membrane proteins or regulators of biofilm formation or motility (12–18), suggesting a role for the Omr sRNAs in the switch between a sessile or a planktonic lifestyle, and in the modification of the membrane composition.

Since OmrA and OmrB are both conserved in many enterobacterial genomes, it might be expected that they also display specific functions. Consistent with this, *btuB* gene is preferentially repressed by the OmrA sRNA in comparison to OmrB, which is explained by the direct pairing of the OmrA specific central region to *btuB* mRNA (19). And while transcription of both *omrA* and *omrB* genes is directly activated by OmpR, OmrA synthesis, but not OmrB, also responds to the RpoS sigma factor (20, 21).

In contrast, no specific target or specific regulator has been described at this stage for the OmrB sRNA. Using a genetic screen to address this question, we have now identified the RstA response regulator as a multicopy activator specific of OmrB. RstA is the regulator of the RstB-RstA TCS, activated at acid pH, and important for bacterial virulence (22–25). Only a few direct RstA targets are known to date (22) and we thus further characterized *omrB* transcriptional regulation by RstA. Interestingly, this uncovered a new role for a major target of the RstB-RstA TCS, the *asr-samT (previously asr-ydgU)* operon, in modulating both positively and negatively the activity of this TCS via two independent mechanisms. This work thus revealed that the acid-responsive RstB-RstA TCS, even though its regulon is relatively small, is also subject to feedback control. It further highlights the role of small proteins such as the 27 aminoacids SamT in modulating the activity of this widespread class of bacterial regulators.

## MATERIAL AND METHODS

### General microbiology techniques

Strains used in this study are derivatives of the MG1655 *Escherichia coli* K-12 strain, and were constructed as indicated in Table S1. Plasmids and oligonucleotides are respectively provided in Tables S2 et S3. For strain and plasmid constructions, strains were grown in LB medium supplemented with antibiotics as needed at the following concentrations: ampicillin 150 μg/mL, chloramphenicol 10 μg/mL, kanamycin 25 μg/mL. X-gal was used at a final concentration of 20 μg/mL. The strain NEB5-α F’I_q_ was used as the recipient for plasmid constructions.

For the experiments performed to study RstB-RstA, cells were grown in minimal A medium (K_2_HPO_4_ 10.5 g/L, KH_2_PO_4_ 4.5 g/L, (NH_4_)_2_SO_4_ 1 g/L, sodium citrate.2H_2_0 0.5 g/L and 1 mM MgSO_4_) containing 0.2% glucose as carbon source, unless otherwise indicated. For growth in this medium at acid pH, K_2_HPO_4_ was omitted and the same phosphate concentration was maintained with only KH_2_PO_4_. The minimal M9 medium was used in one experiment (for Fig. 3), it is composed of Na_2_HPO_4_ 6 g/L, KH_2_PO_4_ 3 g/L, NaCl 0.5 g/L, NH_4_Cl 1 g/L and 1 mM MgSO_4_, and contains 0.2% glucose as carbon source. The fluorescence measurements were performed as described (19) in CAG growth medium (which is A medium supplemented with 0.25% casamino acids and 0.5% glycerol).

**Figure 3.**
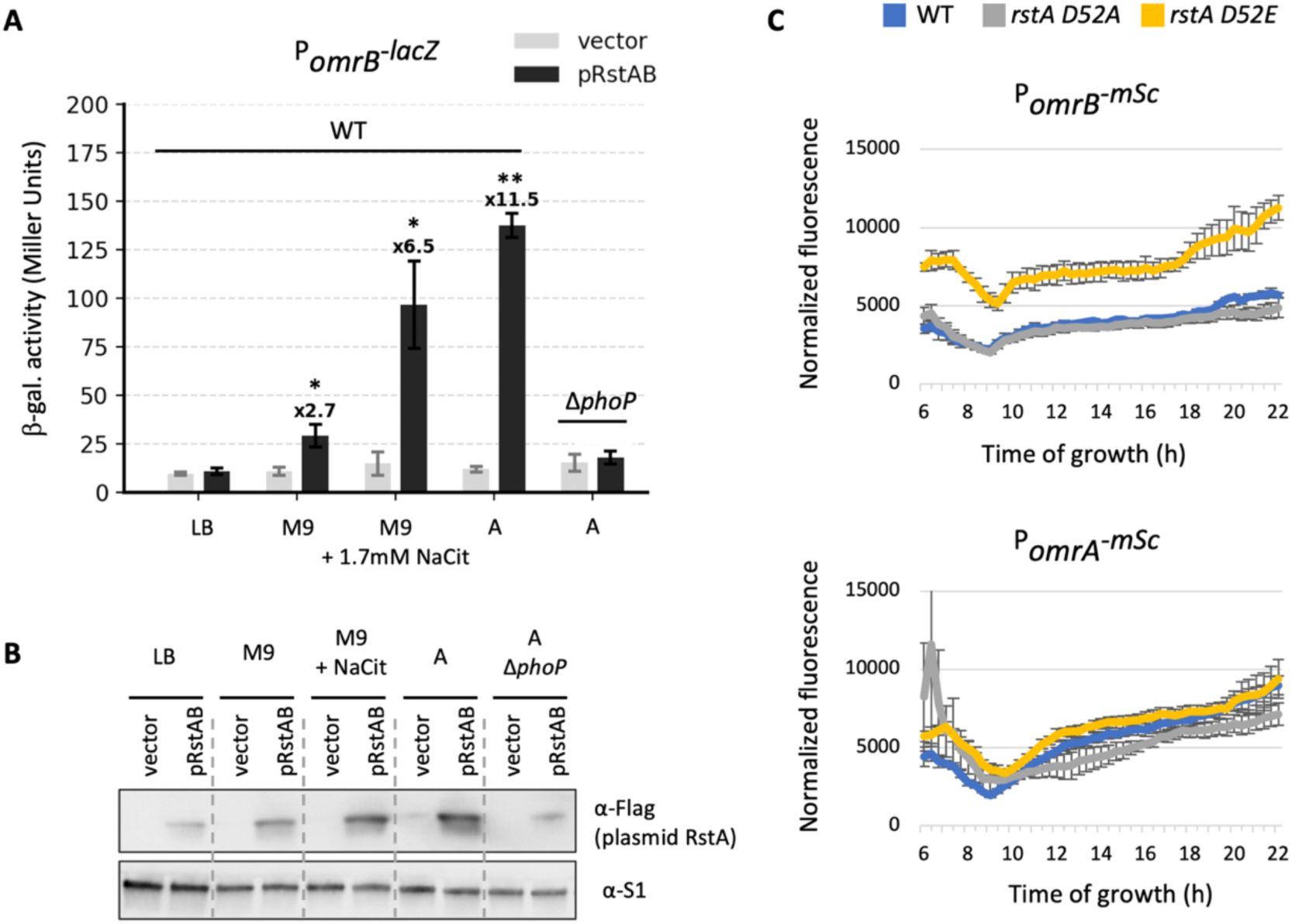
PhoP-dependent transcription of *rstAB* and RstA phosphorylation are required for *omrB* activation by RstB-RstA. (A) Transcriptional activation of *omrB* by pRstAB was measured in exponential phase in the indicated growth media (supplemented with ampicillin) and strain backgrounds. Strains used are JM2110 (WT) and JM2205 (Δ*phoP*). (B) Western blot analysis of the RstA protein made from the pRstAB plasmid, using a derivative of this plasmid with a 3xFlag sequence inserted at the C- terminus of RstA, and detection with an anti-Flag antibody. The levels of the S1 protein were analyzed as a loading control. Strains and growth conditions are as in panel (A). (C) Normalized fluorescence of the P*_omrA_*- and P*_omrB_*-*mSc* reporter fusions in WT or the *rstA* mutants D52A or D52E in minimal A medium at pH 7. The data shown correspond to the average of fluorescence values divided by the absorbance at 600 nm from three independent experiments, with error bars indicating the standard deviations. Strains used in this experiment are JM210, JM287 and JM289 (for P*_omrA_*-*mSc*), and JM211, JM294 and JM296 (for P*_omrB_-mSc*).

### Sequence alignment

For the alignment of the *omrAB* locus, sequences were gathered from the KEGG bank by taking the regions ranging from +50 in *galR* coding sequence to +50 in *aas* coding sequence. Multiple alignment was done with ClustalW using the ClustalW DNA matrix (26). Parameters used were 10 for gap opening penalty and 4 for gap extension. The phylogenetic tree was created using 16S rRNA sequences from each genome, which were also multiple aligned with ClustalW. All genomes used are indicated in Table S4.

### Homolog Identification of Asr and SamT

Genomes were gathered on the basis of complete assemblies. Protein-coding sequences were predicted using Prodigal v2.6.3 (27) or extracted from GFF annotations when available and translated to amino acid sequence. Homologs of the Asr and SamT were then identified with BLASTP (28) using *E. coli* K-12 sequence as an input and a 1e_-5_ e-value threshold. For genomes with positive hits, complementary Hidden Markov Model profile searches were conducted using HMMER v3.3 (29). Results were merged, eliminating duplicates based on genomic coordinates. All genomes used are indicated in Table S5.

### Genetic screen

Electrocompetent cells of strain MG1812 were transformed with the genomic DNA library constructed in the pHDB3 vector (from (30)), and dilutions were plated on minimal A glucose plates containing ampicillin and X-gal. A total of about 7500 colonies were screened, out of which 48 appeared reproducibly bluer than the average. After purification and retransformation of the corresponding plasmids into the initial strain, 14 plasmids that still activated the expression of the P*_omrB_-lacZ* fusion on plates were sequenced.

### β-galactosidase assay

Saturated overnight cultures were diluted 200-fold in fresh medium (500-fold dilution when using LB) and grown to exponential phase. When reaching an optical density of about 0.4 at 600 nm, 0.5 mL of cell culture was mixed with 0.5 ml Z buffer and the β-galactosidase activity was assayed following the Miller protocol (31). The data shown correspond to the average of three independent experiments, and the error bars indicate the standard deviation. Statistical significance of the results was assessed by calculation of the p-value through a heteroscedastic two-tails t test. P-values are represented by an asterisk when the values are significantly different (* for p-values ≤0.05, ** for p-values ≤0.01).

### Fluorescence measurement

Saturated overnight cultures were diluted 200-fold in 200 μL CAG medium in 96-wells plate (Greiner #655090), covered with 50 μL mineral oil, and incubated at 37°C with shaking. Bacterial growth and fluorescence were followed for 16 to 24 hours by measuring the optical density at 600 nm and the fluorescence (excitation at 560±15 nm, emission at 600±15 nm with 580 nm dichroic filter) every 20 minutes on a BMG Labtech Clariostar plus. Results are presented as average fluorescence and standard deviations from three independent replicates. All time points are shown for experiments with only a limited number of samples; otherwise, a single representative time point is presented.

### RNA extraction and northern blot

2 mL of cells grown to exponential phase in minimal A medium were pelleted by centrifugation at 4°C for 2 minutes and resuspended in 650 μL PBS 1X, which were used for RNA extraction using the hot phenol procedure as previously described (12). After ethanol-precipitation, the pellets were resuspended in 15 μL H_2_0. Total RNA was quantified by measuring the absorbance at 260 nm, and a constant amount (4 or 5 μg) were separated on a 6% acrylamide gel. Transfer to nitrocellulose and detection of RNAs using specific biotinylated probes was as in (32). If needed, the same membrane was sequentially probed for different RNAs after boiling 10 minutes in 0.5 % SDS and rinsed with deionized water.

### Western blot

Protein samples preparation and western blotting were performed mostly as described previously (4), with proteins separated on Mini-protean TGX precast gels 4–15% (Biorad) Transfer was done with the Trans- blot Turbo RTA transfer kit Nitrocellulose (Biorad) and detection of the flagged RstA was performed with the Anti-Flag M2 monoclonal antibody- alkaline phosphatase conjugate (Sigma #A9469) following manufacturer’s instructions. The detection of the S1 protein, using an anti-S1 antibody (a gift from K. Nierhaus) at a 1:10 000 dilution overnight in PBS-Tween 0.01% with 2.5% non- fat milk at 4^◦^C, was performed with the Clarity Max Western Substrate (Biorad).

### Purification of RstA-His and OmpR-His proteins

The OmpR-His protein was expressed from the pET15b plasmid derivative (from (33)) and purified as described previously (4). The *rstA* gene was cloned in the pET28 plasmid and the resulting RstA-His protein (with the His-tag at the N-terminus) was purified according to the same procedure. When needed, the proteins were phosphorylated *in vitro* in the presence of acetyl-phosphate following again the same procedure than in (4).

### *In vitro* transcription

Plasmid DNAs derivatives of the pRLG770 plasmid (34) carrying the *asr* (-250+50) or *omrB* (- 104+50) regions relative to the transcription start site, upstream of the 5S rRNA gene, were used as templates. For the transcription reactions, 50 ng of plasmid were mixed with the indicated amount of purified OmpR- or RstA-His and 0.45 u of *E. coli* RNA polymerase holoenzyme (NEB #M0551S) in the presence of 5 mM DTT and 0.5 mg/mL BSA, in TMK buffer (Tris-HCl pH7.6 20 mM, MgCl_2_ 10 mM, KCl 100 mM), and equilibrated at 37°C for 10 minutes. The rNTPs were then added at a final concentration of 0.2 mM for ATP, CTP and GTP, and 10 nM UTP, in a mix containing also 2.5 μCi [α-_32_P] UTP, to initiate transcription, and reaction was incubated at 37°C. After 20 minutes, the reaction was stopped by addition of 1 volume of stop solution (95% formamide, 25 mM EDTA, 0.05% xylene cyanol and bromophenol blue). Transcription products were separated on a 6% sequencing gel, that was dried and imaged with a Typhoon phosphorimager.

### DRaCALA assay of DNA-protein interaction

The interaction between purified RstA-His and either the *omrB* or the *asr* promoter was analyzed in a differential radial capillary action of ligand assay (DRaCALA, (35, 36)). Each promoter was amplified by PCR using the Q5 high-fidelity polymerase and a forward primer that has been 5’-labeled with _32_P prior to the reaction. 15 fmol of the resulting double-strand DNA were mixed with RstA or RstA-P at the indicated protein/DNA ratio, in the presence of Tris-HCl pH7.6 20 mM, KCl 50 mM, EDTA 0.2 mM, MgCl_2_ 1 mM, DTT 1.25 mM and BSA 1.56 μg/mL, and the reactions were incubated at 37°C for 30 minutes. Drops of 4 μL were deposited on a nitrocellulose membrane, dried at room temperature for 10 minutes and the membrane was imaged on a Typhoon phophorimager.

### Bacterial two-hybrid assay

BTH101 cells transformed with the different combinations of pKT25 and pUT18C derivatives were grown overnight in deep-well plates (500 µL LB, Amp 100 µg/mL, Kan 50 µg/mL, IPTG 100 µM, in each well) at 30°C using a thermomixer (1400 rpm) for agitation. A 2.5 µL drop was then spotted on MacConkey agar containing 1% maltose, Amp 100 µg/mL, Kan 50 µg/mL and IPTG 100 µM, let to dry for 5 minutes at room temperature, and incubated at 30°C. Images were taken after 16 h.

## RESULTS

### Promoter regions of *omrA* and *omrB* exhibit specific conservation patterns suggesting differential regulation of these two sRNAs

OmrA and OmrB genes are located adjacent in the *aas-galR* intergenic region and are expressed as individual transcription units with their own promoters (Fig. 1A). This organization is conserved in *Enterobacteriaceae* and aligning the *aas-galR* intergenic region showed conservation both in the *omrA* and *omrB* transcribed sequences, but also in their promoter regions (Fig. 1, B and C, and Fig. S1). More precisely, in addition to the -35 and -10 sequences, four regions in the *omrA* promoter and two in the *omrB* promoter displayed high conservation. Interestingly, and despite their common regulation by the EnvZ-OmpR TCS, these conserved motifs differed between *omrA* and *omrB*, suggesting the existence of specific regulators of either sRNA as well.

This analysis also highlights that while the two sRNAs are present in many species, there are several examples where only one is present in the *aas-galR* region. In such cases, the single sRNA is either OmrB, or a chimeric sequence combining the features of the *omrB* promoter and the *omrA* transcribed region. In other words, the *omrB* promoter is always present, in contrast to the *omrA* promoter or either OmrA or OmrB transcript.

Hence, these data suggest the possibility of a differential regulation of the OmrA and OmrB sRNAs but, to date, no specific OmrB regulator has been identified. This led us to perform a genetic screen in order to unravel OmrB regulators other than the known EnvZ-OmpR TCS.

### Specific activation of *omrB* transcription by the RstB-RstA TCS in multicopy

To search for novel OmrB regulators, we screened for plasmids from a genomic DNA library (from (30)) that would modulate the activity of a reporter P*_omrB_-lacZ* fusion. Because plasmids previously identified in rich medium with a similar approach either carried the *ompR* gene or were epistatic to *envZ-ompR* (12), the present screen was performed in an *ompR* deleted strain, to exclude candidates affecting OmpR synthesis or activity. We also chose to perform this screen on minimal A medium plates with the rationale that regulators others than these identified previously in rich medium might be discovered in these conditions. We isolated 14 plasmids that up-regulated the expression of the P_omrB_-lacZ fusion on plates, and their inserts were sequenced (Fig. 2, A and B).

They contain 8 distinct but possibly overlapping DNA fragments, originating from 4 different regions of *E. coli* genomic DNA. As expected, several of these inserts (4 non-identical ones, present in nine plasmids in total) carried the *ompR* gene encoding the known OmrA/B regulator, showing that OmpR is also an activator of *omrB* in minimal medium. In addition, one plasmid carried the *rstAB* operon, together with the *ydgC* upstream gene, and the *folM* and *tus* adjacent genes, both truncated. Consistent with the screen, this plasmid activates the P*_omrB_-lacZ* fusion in A minimal medium in a Δ*ompR* background. This was observed both on plate and in liquid culture (>33-fold activation in liquid, Fig. 2, B and C), and Northern blot analysis showed that it activated synthesis of the OmrB sRNA as well (Fig. 2D). Furthermore, even though pRstAB was isolated in a Δ*ompR* strain, it also efficiently activates *omrB* in the presence of the chromosomal *ompR* gene (Fig. 2, C and D, similar *omrB* expression in the presence of pRstAB in *ompR_+_* and Δ*ompR* backgrounds). This shows that the effect of the pRstAB on *omrB* transcription is not restricted to conditions where *ompR* is inactivated.

Out of the genes present on this plasmid, the *rstAB* operon is the most likely to explain the effect on *omrB* expression, as it encodes a TCS composed of the RstB sensor kinase and the RstA response regulator. This was more directly tested by inserting two premature stop codons in each of the ORF fully present on the plasmid: *rstA*, *rstB* or *ydgC* (represented by ** in Fig. 2E). This mutation in *rstA* completely abolished the *omrB* activation while the introduction of stop codons in *ydgC* had no effect (Fig. 2E). Interestingly, the introduction of stop codons in *rstB*, or mutation to an *rstB* mutant allele that no longer phosphorylates RstA *in vitro* (*rstB-LA*, (37)) led in contrast to a stronger activation of *omrB* (22-fold in each case, *vs* 11.5 for the pRstAB, Fig. 2E). Overall, these results strongly indicate that the RstA protein is responsible for the effect of the pJ2 plasmid, while RstB is likely involved in modulating RstA activity. The pJ2 plasmid was thus renamed pRstAB. As for the other plasmids isolated in this screen, three of them did not activate *omrB* expression in a liquid assay (pC3, pD1 and pM2, data not shown), while the phenotype associated with pO1 was not always reproducible (Fig. 2B). These four plasmids were not investigated further.

We next tested whether pRstAB affected *omrA* expression as well using both a P*_omrA_- lacZ* transcriptional fusion as a reporter or detection of OmrA sRNA by Northern blot. Interestingly, pRstAB had no significant effect on the expression of the P*_omrA_-lacZ* fusion, in either an *ompR_+_* or Δ*ompR* background (Fig. 2C), and it only modestly increased the OmrA levels in the Δ*ompR* background (Fig. 2D). Hence, RstB-RstA TCS is an OmpR-independent multicopy activator of *omrB* transcription, but not of *omrA*, which represents the first report of conditions leading to the specific activation of OmrB synthesis.

### Activation of *omrB* by pRstAB is dependent on the PhoQ-PhoP TCS

These results establish that *rstAB* is as a multicopy activator of *omrB* in minimal medium A, and we next tested whether this effect was visible in other growth media. For this, we assessed the expression of the P*_omrB_-lacZ* reporter in the presence of pRstAB in another minimal medium (M9) and in a rich medium (LB). The activation was strongly impaired in both media, as pRstAB activated the fusion only 2.7-fold in M9 (compared to 11.5-fold in A medium), and no longer affected *omrB* expression in LB (Fig. 3A).

The transcription of *rstAB* operon is under the positive control of the well-characterized PhoQ-PhoP TCS that is itself activated under low magnesium conditions (28 and references therein). Hence, the differential regulation of *omrB* in these media could be due to the differential activation of PhoQ-PhoP TCS. This is consistent with the presence of 1.7 mM citrate in the A medium that could limit magnesium availability. The role of citrate was tested more directly by adding the same citrate concentration to M9 medium, which increased the activation to 6.5-fold (Fig. 3A). These data suggest that pRstAB activates *omrB* when *rstAB* transcription is sufficiently high, *i.e.* when it is up-regulated by the PhoQ-PhoP TCS, under low magnesium conditions. In agreement with this, the effect of the pRstAB on *omrB* in A medium was fully abolished in a *phoP*-deleted background (Fig. 3A). The levels of RstA produced from the pRstAB plasmid in the different growth media were further analyzed by western blot using a modified pRstAB with a *rstA*-*3xflag* allele (Fig. 3B). This confirmed that the activation of *omrB* by pRstAB increased with the level of the RstA protein made from the plasmid.

In addition to this effect in multicopy, we investigated the activation of *omrB* by the chromosomal *rstAB*. We did not observe any difference when comparing the OmrB levels, or OmrA as a control, in a WT and Δ*rstA* strains, even under acid stress conditions known to activate the RstB-RstA TCS (Fig. S2). We then followed the expression of *omrA* or *omrB* transcriptional fusions with the *mScarlet (mSc)* fluorescent reporter in strains carrying the WT, phospho-null (D52A) or phospho-mimic (D52E) allele of the *rstA* chromosomal copy (Fig. 3C). The fluorescence of the P*_omrB_* fusion was increased ∼2-fold in the D52E mutant compared to the WT or the D52A context, indicating that the phosphorylated form of RstA activates *omrB* transcription. In contrast, the P*_omrA_*-*mSc* fusion was not significantly affected by any of the mutants, confirming the specificity of the *omrB* regulation.

### The activation of *omrB* by RstB-RstA TCS could be indirect

The next question was whether the activation of *omrB* by the RstB-RstA TCS is due to direct binding of the RstA response regulator to the *omrB* promoter. The presence of a site compatible with RstA binding based on the described consensus (22, 39) in the box B2 of the *omrB* promoter suggested a direct interaction. Mutating this site with the T_-75_C and A_-68_G changes in the P*_omrB_-lacZ* fusion abolished activation by pRstAB (Fig. 4A, “mut” change).

**Figure 4.**
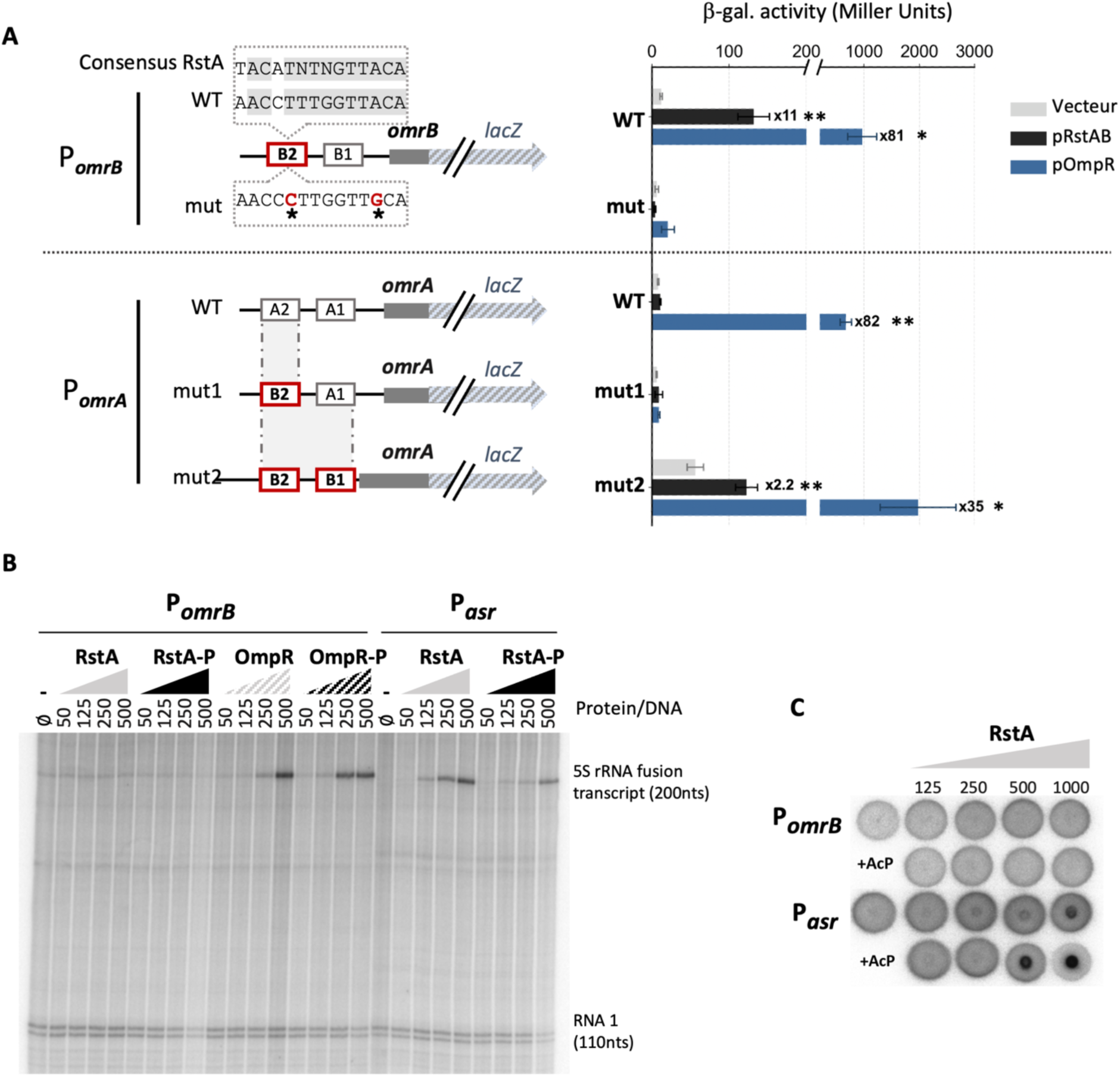
The transcriptional activation of *omrB* by pRstAB is possibly indirect. The activation of transcription from WT or mutant versions of the *omrA* or *omrB* promoters by the pRstAB or pOmpR plasmid was followed by measuring the β-galactosidase activity of the corresponding promoter fusions. Strains used here are JM2110 and JM2132 (for P*_omrB_-lacZ*), and JM2111, JM2130 and JM2136 (for *P_omrA_-lacZ*); they were grown in minimal A glucose medium supplemented with ampicillin. (B) *In vitro* transcription assay on P*_omrB_* or P*_asr_* promoters in the presence of increasing concentration of purified RstA or OmpR protein. The DNA templates are derivatives of the pRLG770 plasmid (34) carrying the indicated promoter driving transcription of an RNA carrying *omrB* or *asr* first 50 nts in fusion with the 5S rRNA sequence (top arrow on the right of the gel). The RNA1 transcript is visible at the bottom of the gel and is used as an internal control for the reaction (bottom arrow). When indicated, the RstA or OmpR proteins were phosphorylated *in vitro* in the presence of acetyl-phosphate prior to the transcription reaction. The numbers above the lanes correspond to the molar ratio of the protein over the plasmid DNA. (C) DRaCALA assay to assess the interaction between the RstA purified protein and the ^32^P-labeled PCR products corresponding to the *omrB* or *asr* promoter regions. The protein/DNA molar ratio is indicated for each reaction. “+AcP” denotes the reactions that were performed using purified RstA phosphorylated in the presence of acetyl-phosphate prior to the incubation with the DNA.

Notably, this change also strongly impaired the activation of *omrB* by pOmpR (3.3-fold vs 81- fold activation in the WT), in agreement with the previously reported OmpR binding-site consensus in this region (12). In addition, introducing this box B2 in the *omrA* promoter in conjunction with box B1 (“mut2” change in Fig. 4A) allowed regulation of OmrA by pRstAB, even if at a lower level than for *omrB* (2.2-fold activation vs 11-fold). Together, these data suggest a direct regulation of *omrB* by RstA, via binding to the B2 site in the *omrB* promoter. Nevertheless, the introduction of the box B2 alone did not promote control by RstA and impaired control by OmpR (Fig. 4A, “mut1” change), showing that box B1 is required as well, which agrees with its high sequence conservation.

To go further, we next performed *in vitro* transcription assays using the *E. coli* RNA polymerase holoenzyme with the sigma70 subunit and a plasmid DNA template in which the region -104 to +50 (relative to the transcription start site) of the *omrB* promoter has been cloned upstream of the 5S rDNA gene. As a control, the promoter of *asr*, a known RstA-target, was cloned in the same vector, with the same organization. Consistent with the known regulation of *omrB* and *asr* genes, their transcription was activated by the addition of purified OmpR or RstA regulator, respectively. In both cases, this activation was visible with or without prior incubation of the response regulator with acetyl-phosphate to phosphorylate it (Fig. 4B). In contrast, the transcription of *omrB* was not affected by RstA in these experiments, even after incubation of the protein with acetyl-phosphate, arguing against a direct regulation of *omrB* by RstA. A possible explanation for this discrepancy is that activation of *omrB* by RstA relies on a sigma factor other than the housekeeping sigma70 present in the transcription reaction. While preliminary tests indicate that the effect of pRstAB is still present in the absence of the FliA, FecI or RpoS alternative sigma factors (Fig. S3), we cannot exclude that one of the other *E. coli* sigma factors is required. We thus decided to assess instead the interaction between RstA and the *omrB* promoter, rather than the transcriptional activation, using a DraCALA assay (35). Briefly, the diffusion on a nitrocellulose membrane of _32_P-labelled PCR products corresponding to either *asr* or *omrB* promoter was analyzed in the presence of increasing concentrations of the RstA protein. Unlike DNA, proteins do not diffuse onto the nitrocellulose membrane and will thus increase the radioactive signal at the center of the spot if there is a protein-DNA interaction. With this assay, we could not detect any interaction between RstA and P*_omrB_*, while we observed a clear binding of RstA on the *asr* promoter, again in the presence or absence of acetyl-phosphate (Fig. 4C). While they do not rule out a possible interaction and subsequent activation of *omrB* expression by RstA under experimental conditions other than the ones used here, these *in vitro* data nonetheless suggest that additional factor(s) could be involved in the RstA-mediated regulation of *omrB*.

### Asr is involved in the positive feedback control of the RstB-RstA TCS

Several hypotheses can be made regarding the identity of such additional factors. We first tested a possible role for three other response regulators possibly related to the RstB-RstA pathway: PhoB, known to regulate the RstA-target *asr* (40); CpxR that regulates the RstA- predicted target *ompF* (41), and RcsB that is important for survival in severe acid stress, a condition in which *asr* is one of the most strongly activated genes (42). Deleting any of these three regulators had no effect on the *omrB* activation by the pRstAB plasmid (Fig. S4). We next reasoned that the RstA target(s) themselves could mediate the effect of pRstAB on *omrB in vivo*. To address this possibility, we analyzed the control of *omrB* by pRstAB in strains deleted for either of the best characterized RstA-targets, *asr* and *csgD* (22). Again, the deletion of *csgD* had no effect but, interestingly, *omrB* activation was significantly impaired in the Δ*asr* mutant strain (2.8-fold activation compared to 11-fold in the WT strain, Fig. 5A). The same result was obtained when following the OmrB sRNA by Northern-blot (Fig. 5B): the increase in OmrB levels in the presence of the pRstAB plasmid is lower in the Δ*asr* background compared to the WT strain.

**Figure 5.**
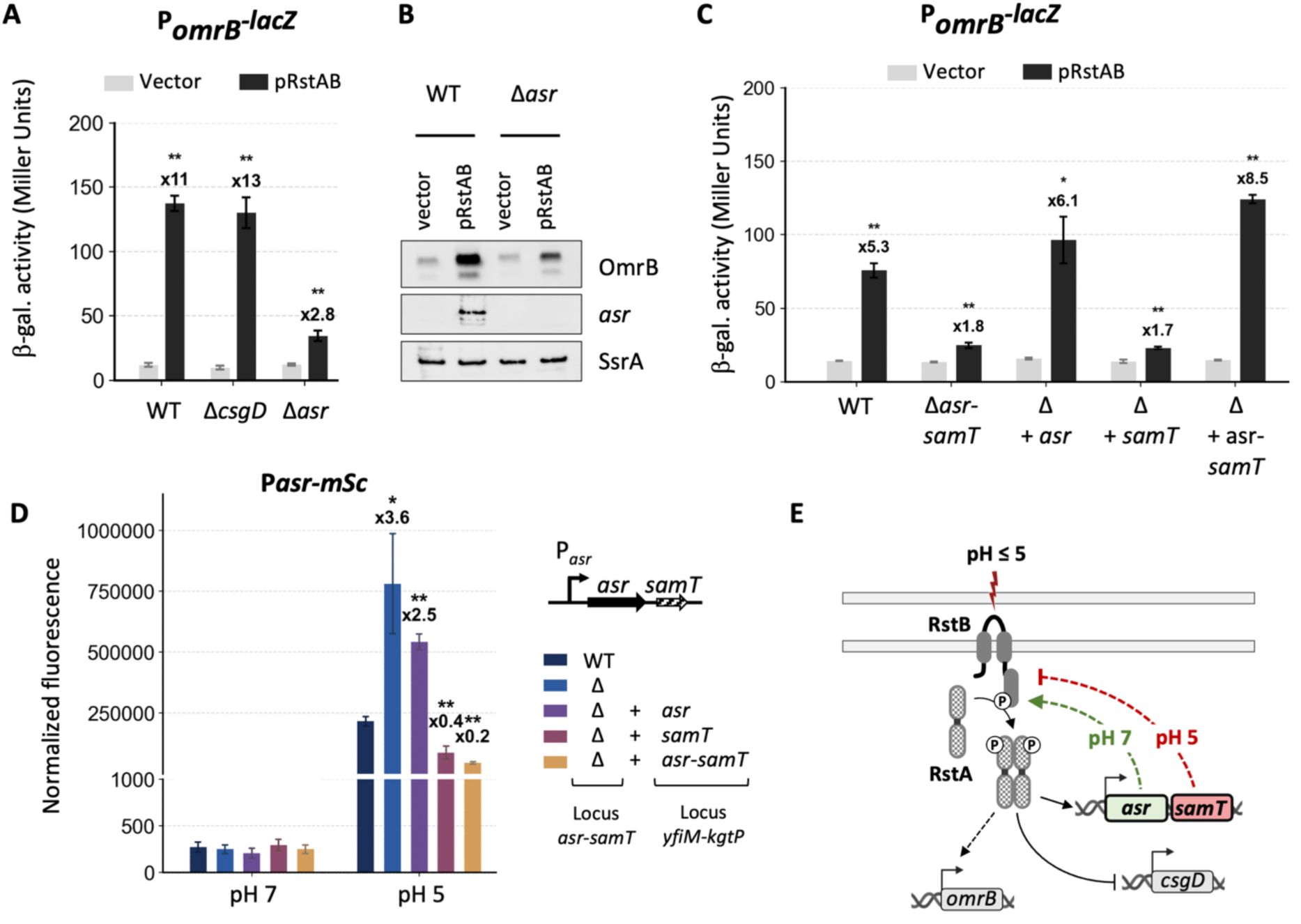
The *asr-samT* operon is involved in the feedback control of the RstB-RstA TCS. (A) Activation of *omrB* transcription by pRstAB in WT, Δ*csgD* or Δ*asr* strains (Keio mutant alleles), as measured by the β-galactosidase activity of the P*_omrB_-lacZ* fusion in minimal A glucose medium, supplemented with ampicillin. Strains used here are JM2110, JM100 and JM161, respectively. (B) OmrB and *asr* RNA levels were assessed by Northern blot from the same cultures as used in (A) in the WT and Δ*asr* strains. (C) The activation of P*_omrB_-lacZ* by pRstAB was analyzed in WT, Δ*asr-samT*, and Δ*asr-samT* cells expressing *asr*, *samT*, or *asr-samT* from the *yfiM-kgtP* locus, as schematized in panel D and Fig. S5. Strains used in this experiment are JM348 (WT), JM349 (Δ), JM350 (Δ+*asr-samT*), JM351 (Δ+*asr*) and JM352 (Δ+*ydgU*). (D) Complementation of the Δ*asr* effect on P*_asr_-mSc* fusion by *asr*, *samT* or the *asr-samT* operon, located *in trans* at the *yfiM-kgtP* locus. Experiment was carried out as in (C) using strains JM116, JM337, JM340, JM342 and JM338, respectively, grown in CAG medium at pH 7 or equilibrated at pH 5 with phosphate buffer. Cells were grown for 17 hours and the data are shown for the time-point closest to an optical density of 0.15. Raw data for panel D are provided in Fig. S7. (E) Summary of the dual effect of the *asr- samT* operon on RstB-RstA signaling at pH 5 or pH 7.

The Δ*asr* used here comes from the Keio collection, and consists in a replacement of the *asr* ORF by a kanamycin resistance cassette (43). Importantly, since the construction of this library, the *samT* (previously *ydgU*) small ORF, encoding a 27 aa peptide, has been identified downstream of, and in operon with, *asr* (44, 8). The *asr* and *samT* coding regions are separated by 93 bp and we cannot exclude that this Δ*asr* mutant also prevents expression of *samT*. Hence, our results indicate that the *asr* and/or the *samT* genes are required for maximal activation of *omrB* by RstA, even though they are not essential for this regulation. Asr is a positively charged periplasmic protein that is important for survival at acid pH and outer membrane integrity, and was found to either promote or prevent protein aggregation depending on the substrates (45, 46). As *asr*, *samT* is expressed under acid stress or low Mg2+ environment, *i.e.* conditions that activate RstB-RstA TCS activity or transcription by PhoQ-PhoP, respectively (42, 8). Acid pH also strongly increases *samT* translation, consistent with the canonical Shine-Dalgarno motif present upstream of *samT* ORF (42). To date, however, no function of the SamT peptide, likely localized at the inner membrane as it carries a predicted transmembrane domain (8), has been reported.

Since Asr is a periplasmic protein and SamT a predicted inner membrane peptide, neither of them is likely to directly participate to the transcriptional activation of *omrB* by RstA; this is also consistent with this control not being fully abolished in the Δ*asr* background (Fig. 5, A and B). To address which part of the *asr-samT* operon is responsible for these effects of the Δ*asr* mutant, we then turned to complementation studies. For this, we expressed the complete *asr-samT* operon or derivatives lacking either *asr* or *samT* gene from an heterologous locus (the *yfiM-kgtP* intergenic region), in a background deleted of the WT *asr- samT* operon (Fig. 5C and S5). The results show that the *asr-samT* operon or *asr* alone are sufficient to restore activation of P*_omrB_-lacZ* fusion by pRstAB to the WT level, while no complementation was observed with *samT* alone. Hence, the *asr* gene promotes the activation of *omrB* transcription by the pRstAB plasmid.

### SamT mediates negative feedback on RstB-RstA at acid pH

Together, these data led us to suspect that the *asr-samT* operon could be involved in a feedback regulation of the RstB-RstA TCS as has already been observed for other TCS-targets, such as MgrB for the PhoQ-PhoP and CpxP for CpxA-CpxR TCSs (10, 47). To investigate this in more detail, we decided to assess the effect of the *asr-samT* operon on RstB-RstA under conditions of this TCS activation, *i.e.* acid pH. As a reporter of RstB-RstA activity, we switched for *asr* promoter, the only positive and direct RstA-target described so far (22). Importantly, this allowed us to work with physiological concentrations of the RstB-RstA TCS rather than using the multicopy plasmid pRstAB.

We thus measured the induction at acid pH of a P*_asr_-mSc* fluorescent fusion. Under these conditions, expression of this fusion is fully dependent of the RstB-RstA TCS (Fig. S6), which agrees with previously published data (22). Surprisingly, when we compared WT and Δ*asr- samT* strains, expression of the RstA-dependent P*_asr_-mSc* was increased in the Δ*asr* strain (3.6- fold increase, Fig. 5D), indicating this time a negative effect of Asr and/or SamT on RstB-RstA. This is in striking contrast with the positive effect of Asr described above. Furthermore, when using the same complementing alleles as in Fig. 5C, we found that either *asr-samT* or *samT* alone, but not *asr*, restored expression of P*_asr_*-*mSc* at the WT level (or even lower) (Fig. 5D and Fig. S7). These data clearly show that the SamT small protein is responsible for the negative feedback on RstB-RstA at acid pH.

In sum, the *asr-samT* operon operates a dual feedback control of the activity of the RstB-RstA TCS: first, a positive one via Asr at neutral pH (Fig. 5C) and, second, a negative one via SamT in acid medium (Fig. 5D); this is summarized in the model of Fig. 5E.

### The conserved SamT peptide inhibits phosphotransfer in the RstB-RstA TCS by direct interaction with the RstB sensor kinase

We next wondered whether this impact of SamT at acid pH was due to an effect on RstB-RstA activity or its synthesis by using different mutants in the *rstAB* chromosomal locus. Since Asr and SamT are periplasmic and inner membrane proteins, respectively, they could signal, directly or indirectly, to the RstB sensor kinase and affect the phosphorylation pathway of RstB-RstA. We therefore used several mutants in which this pathway is abolished: the phospho-null or phospho-mimic version of the regulator as previously (*rstA* D52A or D52E, respectively), a mutant of the sensor kinase that no longer phosphorylates RstA (*rstB-LA*, (37)) or whose gene is interrupted by the introduction of premature stop codons (*rstB\*\**). Strikingly, the feedback control by SamT was lost in all four mutants, as expression of the P*_asr_-mSc* fusion remained unchanged in *asr_+_* and Δ*asr* cells in these backgrounds (Fig. 6A). In contrast, replacing the *rstAB* promoter by a heterologous promoter that is no longer subject to PhoP regulation (here, P_Llac0-1_ in the presence of 100 μM IPTG) did not affect the ability of Asr to control its own expression (Fig. 6A, construction P_lac_-*rstAB*). Together, these results strongly indicate that the *samT* gene exerts its negative feedback regulation at acid pH by modifying the RstB phosphorylation status and/or impairing the phosphotransfer in the RstB-RstA TCS.

**Figure 6.**
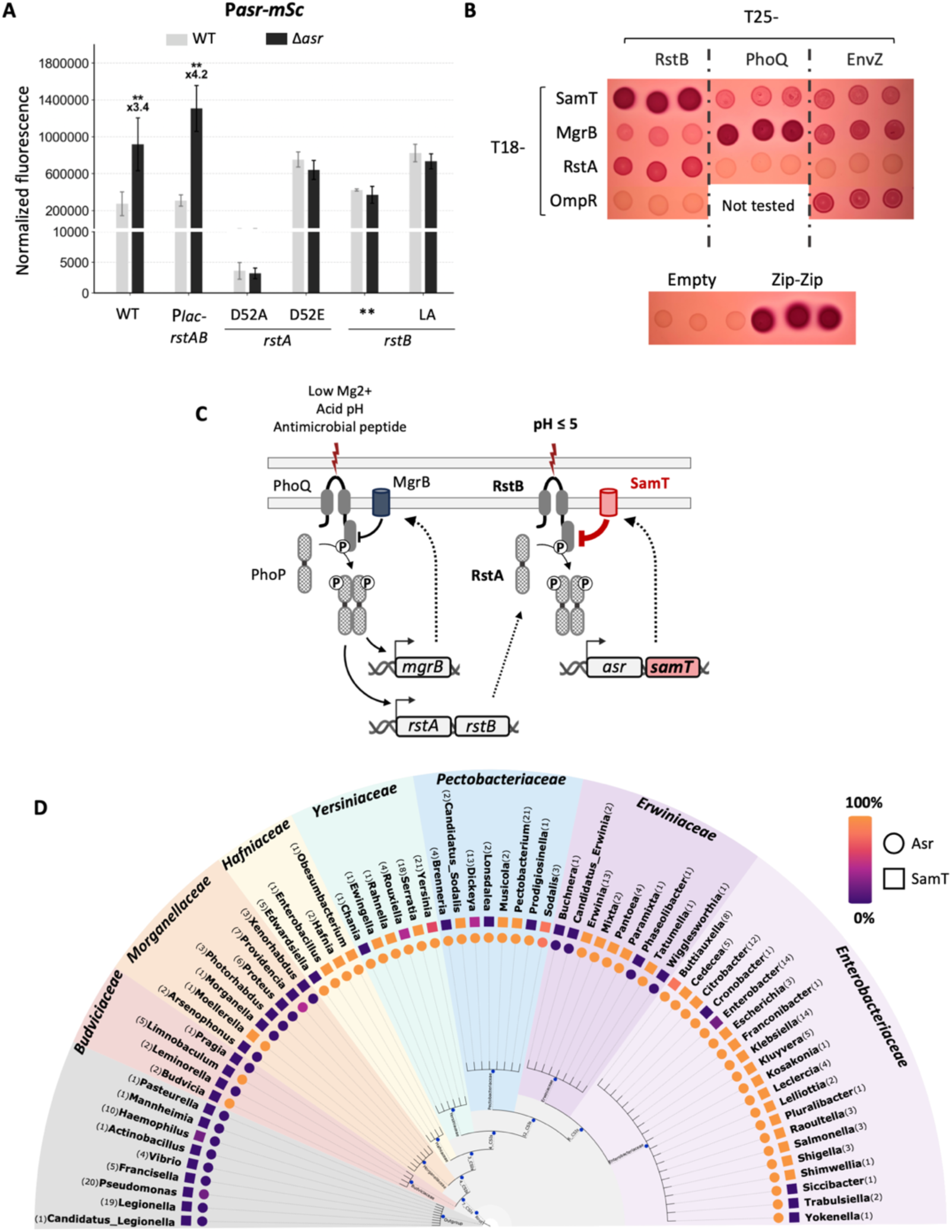
SamT small protein is conserved in y-proteobacteria and directly interacts with the RstB sensor kinase. (A) The effect of *asr-samT* operon on its own transcription was analyzed by following the fluorescence of the P*asr-mSc* fusion in *asr^+^* and the Δ*asr* mutant, in *rstAB^+^* (WT) or the indicated mutants of the *rstAB* chromosomal locus. The data show the average normalized fluorescence values (fluorescence divided by the optical density at 600 nm) from three independent experiments with error bars indicating standard variations. The *asr^+^* and Δ*asr* strains used in this experiment are, respectively, JM209 and JM313 (*rstAB^+^*), JM268 and JM320 (Plac-*rstAB*), JM214 and JM314 (*rstAD52A*), JM215 and JM315 (*rstAD52E*), JM216 and JM316 (*rstB\*\**), and JM217 and JM317 (*rstB-LA*). (B) Bacterial two-hybrid assay was performed by spotting overnight cultures of the BTH101 strain transformed by two plasmids expressing the indicated T18- and T25- fusion proteins on a MacConkey-maltose plate supplemented with ampicillin, kanamycin and IPTG. Plate was incubated at 30°C overnight. (C) Model of the negative feedback of RstB-RstA exerted by SamT at pH 5, in parallel to the negative feedback exerted by MgrB on PhoQ-PhoP (10). (D) Detection of Asr and SamT proteins in y-proteobacteria. The numbers in brackets indicate the number of genomes taken into consideration for determining the conservation of Asr (circles) or SamT (squares) in each genus. The list of genomes used for this analysis is in Table S5.

As this feedback is dependent on the RstB-RstA phosphotransfer, and as SamT is a membrane peptide, we reasoned that SamT could interact with the RstB kinase. This was tested in the bacterial two-hybrid assay relying on the reconstitution of the adenylate cyclase activity (48, 49).

The T18 and T25 fragments of the *Bordetella pertussis* adenylate cyclase catalytic domain were fused to the N-terminal extremity of SamT and the N-terminal extremity of RstB, respectively, and expressed from pUT18C and pKT25 plasmids. We also included MgrB (the modulator of PhoQ-PhoP TCS), and the response regulators RstA and OmpR fused to T18. Other controls were the PhoQ or EnvZ sensor kinases fused to T25 domain, in addition to the T18 and T25 fragments alone (empty plasmids, negative control) or fused to leucine zippers (Zip-Zip, positive control). Reconstitution of the cyclase activity upon interaction of proteins fused to the T18 and T25 fragments was followed by expression of the cAMP/CAP-dependent *mal* genes as visible by the red coloration on McConkey-Maltose plates supplemented with ampicillin, kanamycin and IPTG (48, 49). In this experiment, a very strong interaction was detected between the SamT and RstB proteins, similar to the previously demonstrated MgrB- PhoQ interaction (10) and to the positive control (Fig. 6B). Of note, weaker interactions were also visible in this experiment between the SamT and PhoQ or EnvZ kinases, or between MgrB and RstB or EnvZ. Regarding the sensor kinase-response regulator pairs, specific RstB-RstA and EnvZ-OmpR interactions were detected, as expected, but lead to a weaker induction of the *mal* genes than the RstB-SamT or MgrB-PhoQ interactions in this assay (Fig. 6B).

Together, these data point to a feedback circuit in which the SamT peptide interacts with the RstB sensor kinase and thereby limits the activation of the RstB-RstA phosphocascade at acid pH (Fig. 6C).

Finally, these opposite effects of *asr* and *samT* led us to analyze their conservation in γ- proteobacteria. This search revealed that, while Asr is in general more conserved, SamT is nonetheless present in most genomes that include an *asr* gene (Fig. 6D), raising the possibility that the function of this small protein in the RstB-RstA TCS signaling could be conserved beyond *E. coli*.

## DISCUSSION

Starting from the identification of RstB-RstA TCS as a multicopy activator of the transcription of OmrB sRNA, this work unexpectedly revealed a dual role of the *asr-samT* operon in the feedback control of this TCS. The *asr* gene encodes a periplasmic protein important for response to acid stress, and reinforces the activation of *omrB* expression by pRstAB, at least under neutral pH conditions. In contrast, the SamT small membrane protein mediates negative feedback in the RstB-RstA signaling under acid conditions.

### An OmrB-specific and direct activation by RstA?

Our results did not allow us to discriminate between a direct or indirect transcriptional activation of *omrB* by RstA (Fig. 4). While the effect of mutations in the RstA consensus binding-site present in the conserved B2 box of P_omrB_ is fully consistent with a direct regulation, we were unable to detect either an interaction between RstA and P*_omrB_* or a transcriptional activation in *in vitro* assays. One can hypothesize that RstA binding to P_omrB_ requires additional factor(s), for instance a ligand, or post-translational modifications of RstA or epigenetic modifications of the P_omrB_ promoter. Furthermore, the exchange of boxes from the *omrB* to the *omrA* promoter indicates a role for the highly conserved B1 box in the regulation by RstA. This conserved sequence could serve as a binding-site for another transcription factor, possibly acting in coordination with RstA at the *omrB* promoter. Among the putative factors that could be involved in RstA binding to P*_omrB_*, we can hypothesize a role for *dctR*, encoding a transcriptional regulator and found on a plasmid activating *omrB* in minimal solid medium (Fig. 2, A and B). It would also be interesting to test whether the *ydgC* gene, whose divergent orientation from *rstA* is conserved, could play any role in this regulation. At this stage, we cannot exclude that RstA does not bind *omrB* promoter but acts indirectly to regulate its transcription. This would be in agreement with a recent large scale mapping of transcriptional regulators’ binding-sites by ChIP-seq that did not detect RstA binding to *omrB* promoter (50).

Regardless of the activation mechanism, these data clearly show that the *omrA* and *omrB* promoters can integrate distinct environmental signals. While this has already been proposed for *omrA* via its regulation by RpoS (21, 20), this is the first report, as far as we know, that *omrB* transcription also responds to specific conditions. It is furthermore intriguing that RstA, that is evolutionarily close to the OmpR response regulator (51), is found as an activator of *omrB* as well. In addition to the regulation of the *csgD* target, this provides another example of a partially overlapping regulon between these two response regulators. Because OmrB (together with OmrA) represses *ompR-envZ* expression, it furthermore suggests that RstA activation of *omrB* would in turn indirectly limit the synthesis of EnvZ-OmpR TCS, via the OmrB sRNA. This is reminiscent of OmpR-MicF-CpxR circuit where the transcriptional activation of

MicF by OmpR would lead to lower expression of the *cpxR* gene (52, 53). It will be interesting to determine in the future if and how these sRNA-based circuits participate to the TCS signaling, *e.g.* by limiting the cross-talk, especially between evolutionary close TCSs.

### Small proteins and TCSs

It is interesting to note that modulator proteins, and especially small proteins, are another class of factors repeatedly found in TCS circuits. As in the case of sRNAs, they can connect two different TCSs. For instance, the 65 aa protein SafA connects the EvgS-EvgA TCS with PhoQ-PhoP, since expression of *safA* is activated by EvgS-EvgA and SafA promotes PhoQ kinase activity (9). Note that SafA has thus a positive role on PhoQ and mediates the indirect activation of PhoQ-PhoP in response to the activation of EvgS-EvgA. This is in contrast with the sRNAs mentioned above that, in response to the activation of a given TCS, mediate the down- regulation of another TCS. Another major difference is that SafA directly interacts with the PhoQ protein to modulate its activity while the sRNAs target the TCS-encoding mRNAs and thereby regulate expression of the TCS genes.

Other small proteins are involved in the feedback control of TCSs. A first example was reported with the 47 aa *E. coli* MgrB inner membrane protein that inhibits PhoQ-PhoP signaling via a direct interaction with PhoQ. This results in a negative feedback as *mgrB* is directly activated by the PhoQ-PhoP TCS (10). In *Salmonella enterica*, the 34 aa UgtS small protein also mediates a negative feedback on PhoQ-PhoP signaling, by antagonizing the action of the UgtL protein as an activator of the PhoQ-PhoP TCS. Interestingly, *ugtS* and *ugtL* are encoded as an operon whose different mRNA isoforms originating from two distinct promoters dictate the UgtS/UgtL ratio (11).

The results reported here display similarities with these examples of feedback control. As in the case of MgrB, the SamT small membrane protein directly interacts with the kinase of the TCS that controls its own transcription, thereby limiting signaling within the TCS. And as for *ugtS-ugtL*, the *asr-samT* operon encodes both the positive and negative regulators of the activity of the TCS controlling its transcription. Nevertheless, regulation by *asr*-*samT* occurs via a distinct mode of action compared to the *ugtS-ugtL* operon. While UgtS impairs the positive effect of UgtL on PhoQ (11), our complementation results clearly show that Asr and SamT act independently to activate or repress RstB-RstA signaling, respectively.

In contrast to SamT and its interaction with RstB, the mechanism by which the *asr* gene promotes RstB-RstA activation is still unresolved. Several hypotheses can be made in this regard. For instance, Asr could interact with the periplasmic domain of RstB to impact its kinase or phosphatase activity. Alternatively, it could interact with the PhoQ periplasmic domain, and thereby indirectly affect *rstAB* transcription. Other than via a direct interaction with RstB or PhoQ, Asr could also signal to these sensor kinases by resolving the aggregation of periplasmic proteins thanks to its reported chaperone activity (46).

In the course of this study, the use of *rstB* mutants confirmed the role of the RstB-RstA TCS in the activation of *omrB* (Fig. 2E), and in the feedback exerted by the *asr-samT* operon (Fig. 6A), although with results that unexpectedly differed from those obtained with *rstA* mutants. The *rstB*\*\* null variant as well the *rstB-LA* allele that is defective in phosphotransfer to RstA both led to an hyperactivation of *omrB* transcription when introduced in pRstAB, while the *rstA*\*\* mutant abolished activation, as expected. This suggests that, in minimal A medium at pH 7, the RstB sensor kinase exists in the phosphatase mode and inhibits RstA activity. Even though RstA phosphorylation was apparently not required for the interaction and regulation of the *asr* promoter *in vitro* (Fig. 4, B and C), the *in vivo* activation of *omrB* transcription by RstAD52E, but not by the WT or D52A mutant (Fig. 2C), strongly suggests that the phosphorylated form of RstA is responsible for *omrB* regulation. Furthermore, surprising results were obtained as well when assessing the effect of the same *rstB*\*\* and *rstB-LA* mutants on the chromosome at pH 5: in this condition, they abolished the feedback by *asr- samT*, but they did not impair (and even activated) the expression of the P*_asr_*-*mSc* fusion. Whether RstA could be efficiently phosphorylated in the absence of a functional RstB, which could explain these results, will need to be further investigated.

Overall, this study is a further illustration of the role played by modulator proteins and sRNAs in TCS signaling. The recognition of new actors impacting RstB-RstA suggests exquisite regulation in the establishment of the bacterial response to acid stress via this TCS. In particular, it will be interesting to precisely characterize how the dual feedback of RstB-RstA reported here translates in terms of response to different stimuli of various intensities, connections with other TCSs or kinetics of response.

## FUNDING

This project has received funding from the European Research Council (ERC) under the European Union’s Horizon 2020 research and innovation program (Grant agreement No. 818750). Research in the UMR8261 is supported by the CNRS and the “Initiative d’Excellence” program from the French State (Grant “Dynamo”, ANR-11-LABX-0011). The EURGene mentorship program supported the fourth year of PhD of JMdF.

## ACKNOWLEDGMENTS

We thank Nathalie Dautin (IBPC, Paris) for providing the BTH101 strain, and the pKT25 and pUT18C plasmids as well as their derivatives carrying the leucine zippers. We are grateful to Mark Goulian for the plasmids pKT25-EnvZ, pKT25-PhoQ, pKT25-RstB and pUT18C-MgrB, and to Linda Kenney for the pOmpR-His plasmid. We thank Carine Chagneau for advice with the purification of His-tagged proteins, Eliane Hajnsdorf and Jackie Plumbridge for advice on the *in vitro* transcription assays, Jonathan Jagodnik for advice on the Dracala assays. We are grateful to Jackie Plumbridge and Jonathan Jagodnik for critical reading of the manuscript, and to all members of the group for discussions.

## Notes

### Competing Interest Statement

The authors have declared no competing interest.

