## Supplementary files for "A novel peptide modulator of a two-component system revealed by the specific activation of a small RNA in *Enterobacteriaceae*"

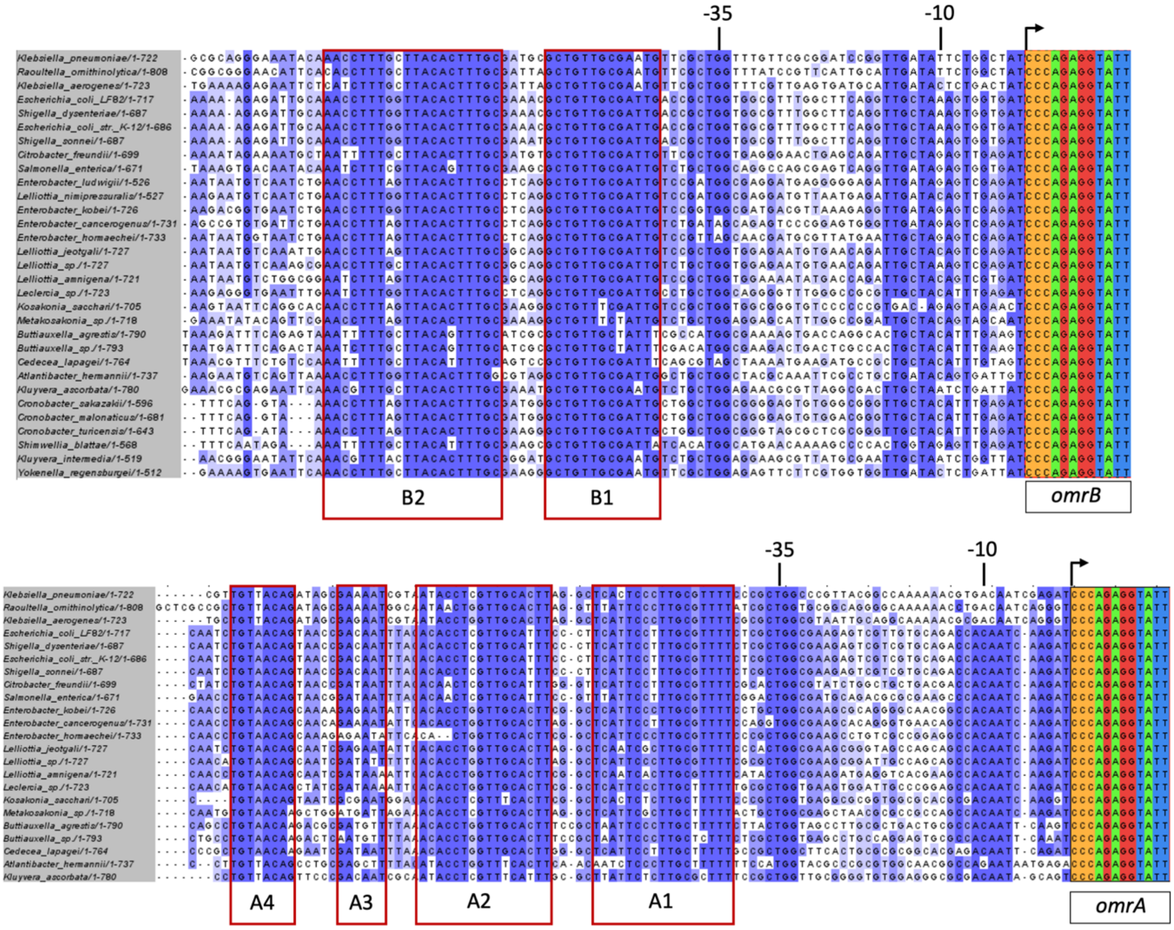

**Figure S1. Alignment of the promoter regions (from -104 to +8, relative to the transcription start site) of *omrA* and *omrB* in *Enterobacteriaceae*.** The conserved boxes A1 to A4 and B1 to B2 are indicated, in addition to the -10 and -35 sequences.

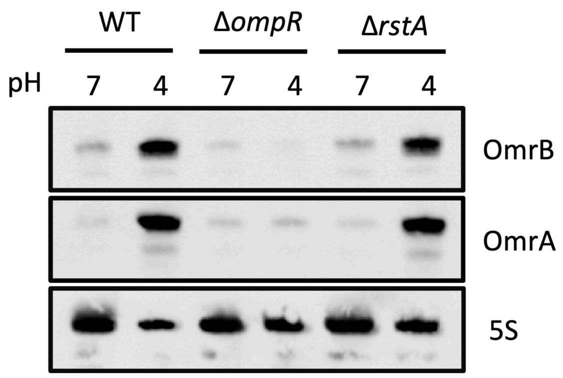

**Figure S2. Northern blot analysis of OmrA and OmrB sRNAs following a one-hour shift at acid pH (or at pH 7 as a control) in WT, Δ*ompR* or Δ*rstA* cells.**

Northern blot analysis of OmrA and OmrB sRNAs, using total RNA extracted from strains JM2110 (WT), JM2118 (Δ*ompR)* and JM2128 (Δ*rstA*). Cells were grown in minimal A medium to an optical density of 0.2 at 600 nm, cultures were then split and centrifuged, and cells were resuspended, either in the same medium or a derivative equilibrated at pH 4 with HCl. Total RNA was extracted after one hour growth in this new medium.

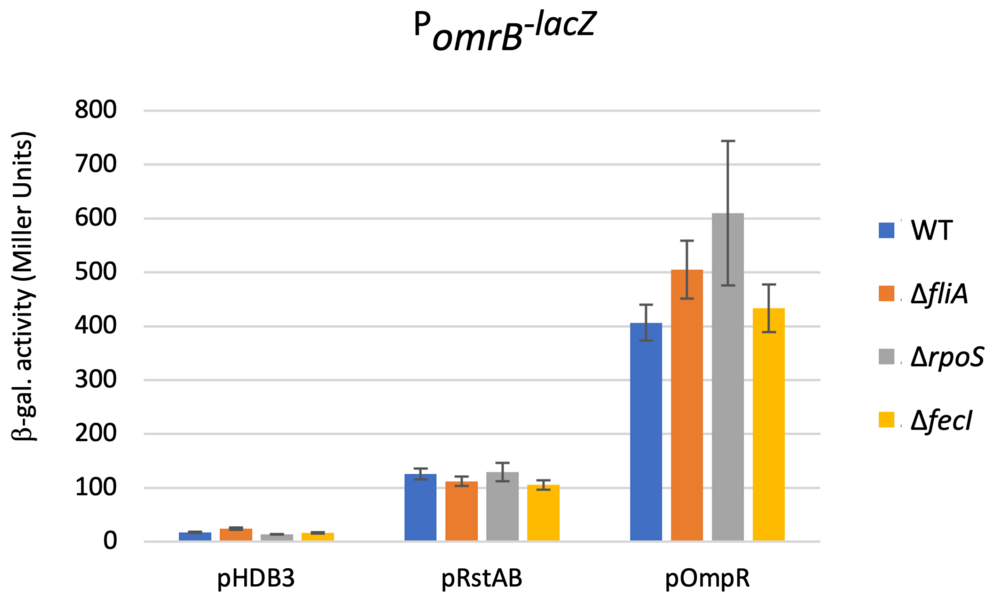

**Figure S3. Activation of the $P_{omrB}$-*lacZ* fusion by the pRstAB and pOmpR plasmids in WT cells or mutants of the FliA, RpoS or FecI alternative sigma factors.**
The *omrB* transcriptional activation by pRstAB was measured in exponential phase in CAG growth medium (supplemented with ampicillin) and strain backgrounds. Strains used are JM2110 (WT), JM122 (Δ*fliA*), JM124 (Δ*rpoS*), JM159 (Δ*fecI*).

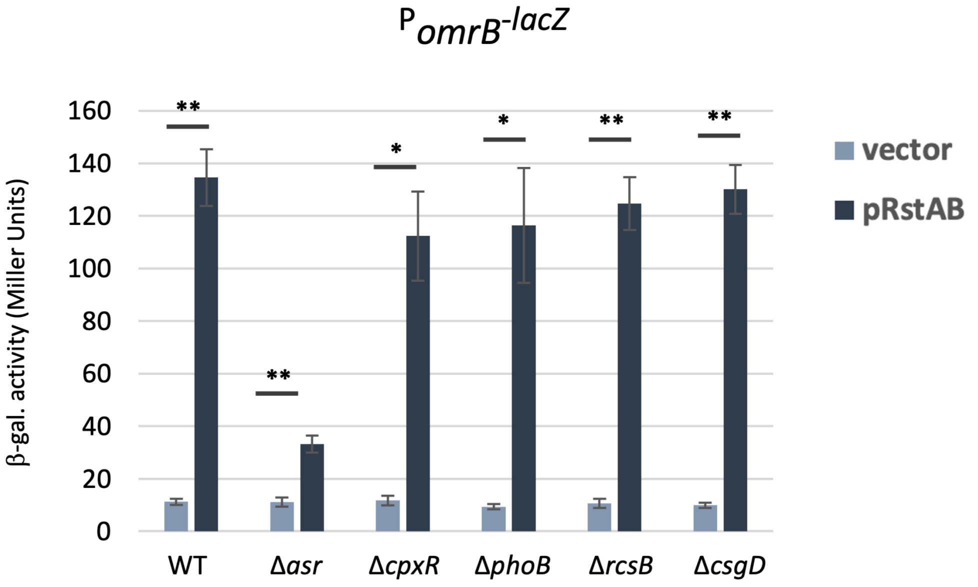

**Figure S4. The activation of *omrB* transcription by pRstAB is independent of *cpxR*, *phoB*, *rcsB* or *csgD*, but is reduced in the absence of *asr*.**
The *omrB* transcriptional activation by pRstAB was measured in exponential phase in minimal A medium (supplemented with ampicillin) and strain backgrounds. Strains used are JM2110 (wt), JM100 (Δ*asr*), JM158 (Δ*cpxR*), JM159 (Δ*phoB*), JM160 (Δ*rcsB*), JM161 (Δ*csgD*).

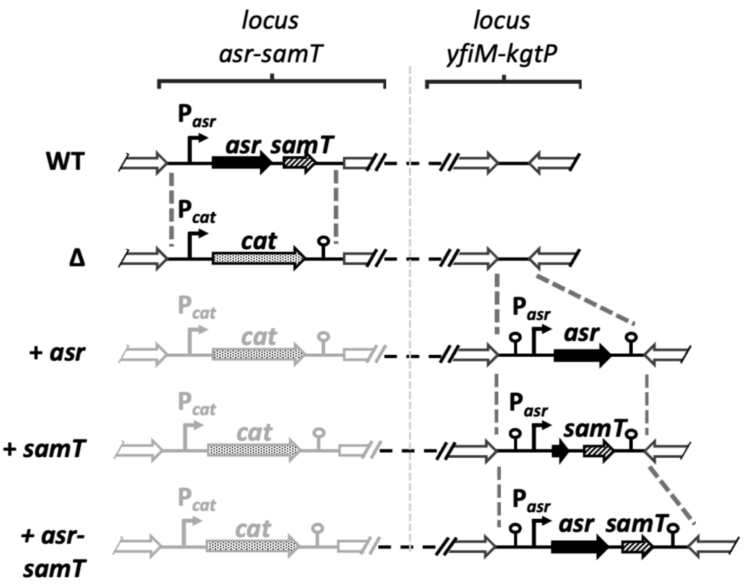

**Figure S5. Scheme of alleles constructed for the functional complementation assays used in Fig. 5C and 5D**.

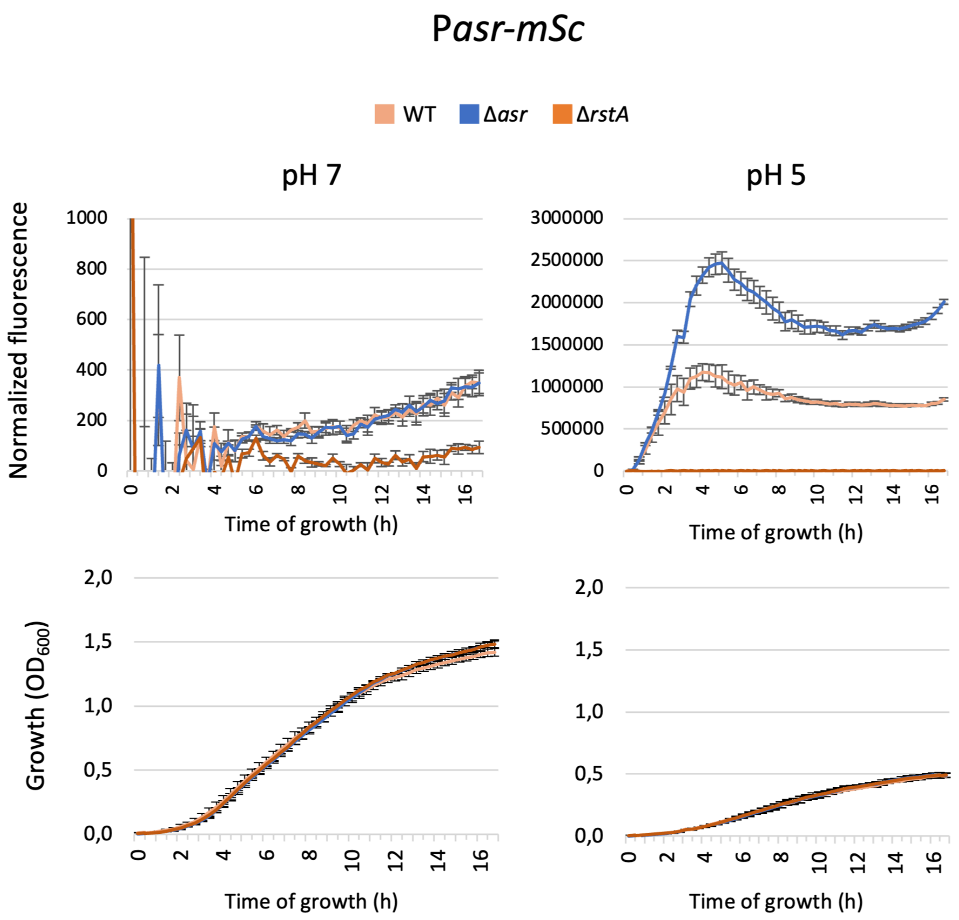

**Figure S6. RstA is essential for transcription from the P$_{asr}$ promoter at acid pH.**
The bacterial growth and fluorescence of WT (JM2210), Δ*asr* (JM142) and Δ*rstA* (JM52) strains carrying a P$_{asr}$-*mSc* transcriptional fusion, were followed in CAG medium at pH 7 or pH 5 for 17 hours.

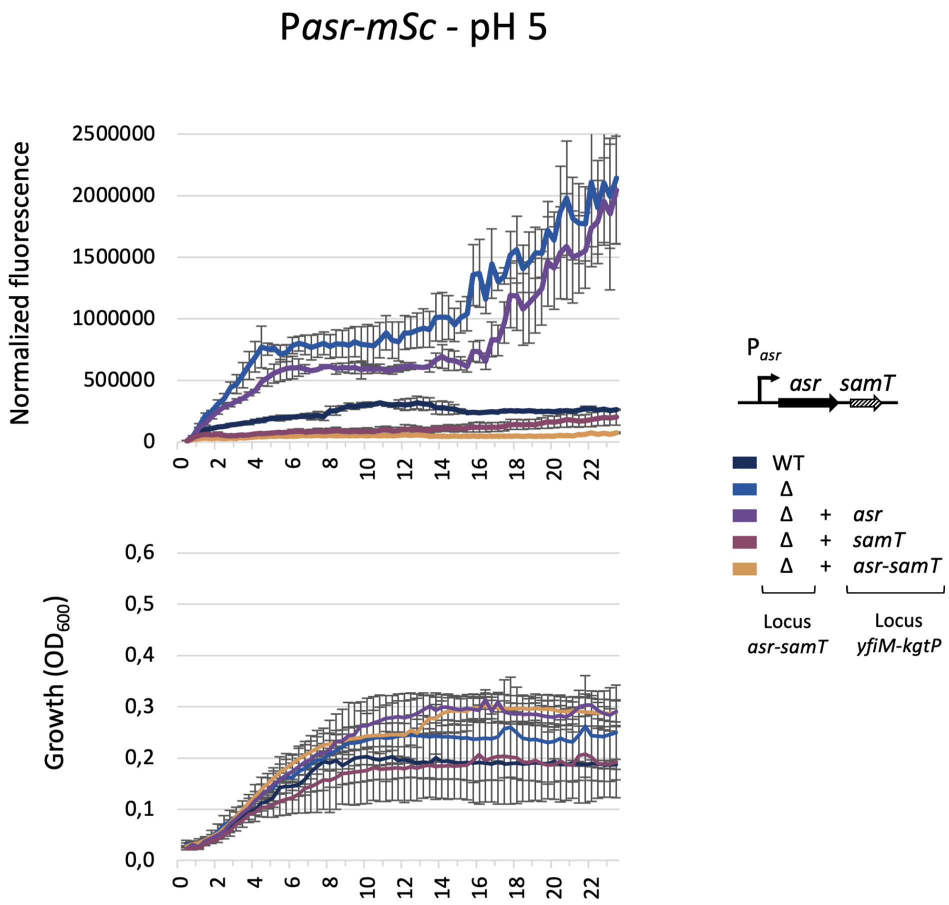

**Figure S7. Raw data for Fig. 5D.**
The growth and normalized fluorescence values are shown for 23 hours of growth. See Fig. 5D for details.

Table S1. Strains used in this study

| **Name** | **Relevant characteristics** | **Source** |
|---|---|---|
| MG1655 | Reference wild-type *Escherichia coli* strain for this study | From F. Blattner's lab |
| DJ480 | MG1655 Δ*lacX74* | D. Jin, NIH |
| DJ624 | MG1655 Δ*lacX74 mal*::*lacI$^q$* | D. Jin, NIH |
| BTH101 | F$^-$, *cya*-99, *araD*139, *galE*15, *galK*16, *rpsL*1 (Str$^R$), *hsdR2*, *mcrA1, mcrB1, relA1* | Euromedex |
| OK510 | DJ624 *argG*-[TT1-PLtetO-1(no -10)-*sacB-cat-'mScarlet*(no ATG)- FRT-*nptII*-FRT-TT2]-*yhbX,* mini-λ tet$^R$ | (1) |
| OK523 | DJ624 *ΔargG*::FRT-*nptII*-FRT | (1) |
| MG1035 | DJ480 Δ*ompR::cat* | (2) |
| MG1432 | DJ624 mini-λ tet | Lab stock |
| MG1433 | DJ624 mini-λ-*cat* | Lab stock |
| MG1508 | MG1655 *mal::lacI$^q$* PLtetO-1-*cat-sacB-lacZ*, mini-λ-*tet* | (3) |
| MG1812 | DJ480 P$_{omrB}$-*lacZ* Δ*ompR::cat* | (4) |
| sML214 | DJ624 *yfiM_TT1_P$_{tet}$_TIR2-flag-lacZ$_{+3+174}$-mScarlet_FRT_Zeo_FRT_TT2_kgtP* | (5) |
| NM300 | DJ480 mini-λ tet | Nadim Majdalani (NIH) |
| JM52 | DJ624 P$_{asr(-250+31)}$-*mSc* Δ*rstA::kan* | This study; JM2210+ P1 (JW1600 from the Keio collection) |
| JM100 | MG1655 *mal::lacI$^q$* P$_{omrB(-104+8)}$-*lacZ* Δ*asr::kan* | This study; JM2110+ P1 (JW5826 from the Keio collection) |
| JM109 | MG1432 Δ*rstAB ::ccdB-cat* | This study; recombineering in MG1432 |
| JM112 | DJ624 *rstA-3xFlag* | This study; recombineering in JM109 |
| JM113 | DJ624 *rstAD52A* | This study; recombineering in JM109 |
| JM115 | DJ624 *rstAD52E* | This study; recombineering in JM109 |
| JM116 | DJ624 P$_{asr(-250+31)}$-*mSc*- FRT-*nptII*-FRT | This study; DJ624 + P1 (JM2134) |
| JM122 | MG1655 *mal::lacI$^q$* P$_{omrB(-104+8)}$-*lacZ* Δ*fliA::kan* | This study; JM2110+ P1 (JW1907 from the Keio collection) |
| JM142 | DJ624 P$_{asr(-250+31)}$-*mSc* Δ*asr::kan* | This study; JM2210+ P1 (JW5826 from the Keio collection) |
| JM158 | MG1655 *mal::lacI$^q$* P$_{omrB(-104+8)}$-*lacZ* Δ*cpxR ::kan* | This study; JM2110+ P1 (JW3883 from the Keio collection) |
| JM159 | MG1655 *mal::lacI$^q$* P$_{omrB(-104+8)}$-*lacZ* Δ*phoB ::kan* | This study; JM2110+ P1 (JW0389 from the Keio collection) |

| | | |
|---|---|---|
| JM160 | MG1655 *mal::*$lacI^q$ $P_{omrB(-104+8)}$*-lacZ* $\Delta$*rcsB ::kan* | This study; JM2110+ P1 (JW2205 from the Keio collection) |
| JM161 | MG1655 *mal::*$lacI^q$ $P_{omrB(-104+8)}$*-lacZ* $\Delta$*csgD ::kan* | This study; JM2110+ P1 (JW1023 from the Keio collection) |
| JM175 | DJ624 *rstB*** | This study; recombineering in JM109 |
| JM177 | DJ624 *rstB-LA (rstB Y232L R233A)* | This study; recombineering in JM109 |
| JM179 | DJ480 $\Delta$*manA ::kan* | This study; DJ480 + P1 (JW1603 from the Keio collection) |
| JM182 | DJ480 *rstA-3xFlag* | This study; JM179 + P1 (JM112) |
| JM183 | DJ480 *rstAD52A* | This study; JM179 + P1 (JM113) |
| JM184 | *DJ480 rstAD52E* | This study; JM179 + P1 (JM115) |
| JM185 | *DJ480 rstB*** | This study; JM179 + P1 (JM175) |
| JM186 | *DJ480 rstB-LA* | This study; JM179 + P1 (JM177) |
| JM209 | DJ480 $P_{asr(-250+31)}$*-mSc-* FRT-*nptII*-FRT | This study; DJ480 + P1 (JM2134) |
| JM210 | DJ480 $P_{omrA(-104+8)}$*-mSc-* FRT-*nptII*-FRT | This study; DJ480 + P1 (JM2203) |
| JM211 | DJ480 $P_{omrB(-104+8)}$*-mSc-* FRT-*nptII*-FRT | This study; DJ480 + P1 (JM2113) |
| JM214 | DJ480 $P_{asr(-250+31)}$*-mSc-* FRT-*nptII*-FRT *rstAD52A* | This study; JM183 + P1 (JM2134) |
| JM215 | DJ480 $P_{asr(-250+31)}$*-mSc-* FRT-*nptII*-FRT *rstAD52E* | This study; JM184 + P1 (JM2134) |
| JM216 | DJ480 $P_{asr(-250+31)}$*-mSc-* FRT-*nptII*-FRT *rstB*** | This study; JM185 + P1 (JM2134) |
| JM217 | DJ480 $P_{asr(-250+31)}$*-mSc-* FRT-*nptII*-FRT *rstB-LA* | This study; JM186 + P1 (JM2134) |
| JM266 | MG1433 *tet*-$P_{lac}$-*rstAB* | This study; recombineering in MG1433 |
| JM268 | DJ624 $P_{asr(-250+31)}$*-mSc tet*-$P_{lac}$-*rstAB* | This study; JM2210 + P1 (JM266) |
| JM287 | DJ624 $P_{omrA(-104+8)}$*-mSc-* FRT-*nptII*-FRT *rstAD52A* | This study; JM113 + P1 (JM2203) |
| JM289 | DJ624 $P_{omrA(-104+8)}$*-mSc-* FRT-*nptII*-FRT *rstAD52E* | This study; JM115 + P1 (JM2203) |
| JM294 | DJ624 $P_{omrB(-104+8)}$*-mSc-* FRT-*nptII*-FRT *rstAD52A* | This study; JM113 + P1 (JM2113) |
| JM296 | DJ624 $P_{omrB(-104+8)}$*-mSc-* FRT-*nptII*-FRT *rstAD52E* | This study; JM115 + P1 (JM2113) |

| | | |
|---|---|---|
| JM312 | MG1432 Δ*asr::ccdB-cat* | This study; recombineering in MG1432 |
| JM313 | DJ480 $P_{asr(-250+31)}$-*mSc*- FRT-*nptII*-FRT Δ*asr::ccdB-cat* | This study; JM209 + P1 (JM312) |
| JM314 | DJ480 $P_{asr(-250+31)}$-*mSc*- FRT-*nptII*-FRT *rstAD52A* Δ*asr::ccdB-cat* | This study; JM214 + P1 (JM312) |
| JM315 | DJ480 $P_{asr(-250+31)}$-*mSc*- FRT-*nptII*-FRT *rstAD52E* Δ*asr::ccdB-cat* | This study; JM215 + P1 (JM312) |
| JM316 | DJ480 $P_{asr(-250+31)}$-*mSc*- FRT-*nptII*-FRT *rstB*** Δ*asr::ccdB-cat* | This study; JM216 + P1 (JM312) |
| JM317 | DJ480 $P_{asr(-250+31)}$-*mSc*- FRT-*nptII*-FRT *rstB-LA* Δ*asr::ccdB-cat* | This study; JM217 + P1 (JM312) |
| JM320 | DJ624 $P_{asr(-250+31)}$-*mSc tet*-$P_{lac}$-*rstAB* Δ*asr::kan* | This study; JM268 + P1 (JW5826 from the Keio collection) |
| JM327 | DJ624 *kgtP*-$P_{asr}$-*asr-ydgU -yfiM* | This study; CRISPR/Cas9-assisted recombineering in sML214 |
| JM329 | DJ624 *kgtP*-$P_{asr}$-*asr-yfiM* | This study; CRISPR/Cas9-assisted recombineering in sML214 |
| JM331 | DJ624 *kgtP*-$P_{asr}$-Δ*asr-ydgU -yfiM* | This study; CRISPR/Cas9-assisted recombineering in sML214 |
| JM335 | NM300 Δ*asr-ydgU::cat* | This study; recombineering in NM300 |
| JM337 | DJ624 $P_{asr(-250+31)}$-*mSc*- FRT-*nptII*-FRT Δ*asr-ydgU::cat* | This study; JM116 + P1 (JM335) |
| JM338a | DJ624 Δ*asr-ydgU::cat* *kgtP*-$P_{asr}$-*asr-ydgU -yfiM* | This study; JM327 + P1 (JM335) |
| JM338 | DJ624 Δ*asr-ydgU::cat* *kgtP*-$P_{asr}$-*asr-ydgU-yfiM* $P_{asr(-250+31)}$-*mSc*- FRT-*nptII*-FRT | This study; JM338a + P1 (JM2134) |
| JM340a | DJ624 Δ*asr-ydgU::cat* *kgtP*-$P_{asr}$-*asr-yfiM* | This study; JM329 + P1 (JM335) |
| JM340 | DJ624 Δ*asr-ydgU::cat* *kgtP*-$P_{asr}$-*asr-yfiM* $P_{asr(-250+31)}$-*mSc*- FRT-*nptII*-FRT | This study; JM340a + P1 (JM2134) |
| JM342a | DJ624 Δ*asr-ydgU::cat* *kgtP*-$P_{asr}$-Δ*asr-ydgU-yfiM* | This study; JM331 + P1 (JM335) |
| JM342 | DJ624 Δ*asr-ydgU::cat* *kgtP*-$P_{asr}$-Δ*asr-ydgU-yfiM* $P_{asr(-250+31)}$-*mSc*- FRT-*nptII*-FRT | This study; JM342a + P1 (JM2134) |
| JM348 | DJ624 $P_{asr(-250+31)}$-*mSc*- FRT-*nptII*-FRT $P_{omrB(-104+8)}$-*lacZ* | JM116 + P1 (JM2110) |
| JM349 | DJ624 Δ*asr-ydgU::cat* $P_{asr(-250+31)}$-*mSc*-FRT-*nptII*-FRT $P_{omrB(-104+8)}$-*lacZ* | JM337 + P1 (JM2110) |
| JM350 | DJ624 Δ*asr-ydgU::cat* *kgtP*-$P_{asr}$-*asr-ydgU-yfiM* $P_{asr(-250+31)}$-*mSc*- FRT-*nptII*-FRT $P_{omrB(-104+8)}$-*lacZ* | JM338 + P1 (JM2110) |
| JM351 | DJ624 Δ*asr-ydgU::cat* *kgtP*-$P_{asr}$-*asr-yfiM* $P_{asr(-250+31)}$-*mSc*- FRT-*nptII*-FRT $P_{omrB(-104+8)}$-*lacZ* | JM340 + P1 (JM2110) |
| JM352 | DJ624 Δ*asr-ydgU::cat* *kgtP*-$P_{asr}$-Δ*asr-ydgU-yfiM* $P_{asr(-250+31)}$-*mSc*- FRT-*nptII*-FRT $P_{omrB(-104+8)}$-*lacZ* | JM342 + P1 (JM2110) |

| | | |
|---|---|---|
| JM2110 | MG1655 *mal::lacI^q^* $P_{omrB(-104+8)}$-*lacZ* (*omrB* promoter from -104 to +8 relative to transcription start site inserted upstream of *lacZ*) | This study, recombineering in MG1508 |
| JM2111 | MG1655 *mal::lacI^q^* $P_{omrA(-104+8)}$-*lacZ* (*omrA* promoter from -104 to +8 relative to transcription start site inserted upstream of *lacZ*) | This study, recombineering in MG1508 |
| JM2113 | DJ624 $P_{omrB(-104+8)}$-*mSc*- FRT-*nptII*-FRT (*omrB* promoter from -104 to +8 relative to transcription start site inserted upstream of *mSc*) | This study, recombineering in OK510 |
| JM2115 | MG1655 *mal::lacI^q^* $P_{omrA(-104+8)}$-*lacZ* Δ*ompR ::cat* | This study ; JM2111 + P1 (MG1035) |
| JM2118 | MG1655 *mal::lacI^q^* $P_{omrB(-104+8)}$-*lacZ* Δ*ompR ::cat* | This study ; JM2110 + P1 (MG1035) |
| JM2128 | MG1655 *mal::lacI^q^* $P_{omrB(-104+8)}$-*lacZ* Δ*rstA ::kan* | This study ; JM2110 + P1 (JW1600 from the Keio collection) |
| JM2130 | MG1655 *mal::lacI^q^* $P_{omrA(-104+8)}$<u>*mut1*</u>-*lacZ* (*omrA*<u>*mut1*</u> promoter from -104 to +8 relative to transcription start site inserted upstream of *lacZ*) | This study, recombineering in MG1508 |
| JM2132 | MG1655 *mal::lacI^q^* $P_{omrB(-104+8)}$<u>*mut*</u>-*lacZ* (*omrB*<u>*mut*</u> promoter from -104 to +8 relative to transcription start site inserted upstream of *lacZ*) | This study, recombineering in MG1508 |
| JM2134 | DJ624 $P_{asr(-250+31)}$-*mSc*- FRT-*nptII*-FRT (*asr* promoter from -250 to +31 relative to transcription start site inserted upstream of *mSc*) | This study, recombineering in OK510 |
| JM2134b | DJ624 $P_{asr(-250+31)}$-*mSc* (KanS) | This study, nptII cassette flipped using pCp20 |
| JM2136 | MG1655 *mal::lacI^q^* $P_{omrA(-104+8)}$<u>*mut2*</u>-*lacZ* (*omrA*<u>*mut2*</u> promoter from -104 to +8 relative to transcription start site inserted upstream of *lacZ*) | This study, recombineering in MG1508 |
| JM2203 | DJ624 $P_{omrA(-104+8)}$-*mSc*- FRT-*nptII*-FRT (*omrA* promoter from -104 to +8 relative to transcription start site inserted upstream of *mSc*) | This study, recombineering in OK510 |
| JM2205 | MG1655 *mal::lacI^q^* $P_{omrB(-104+8)}$-*lacZ* Δ*phoP::cat-sacB* | This study ; JM2110 + P1 (MG1633) |
| JM2210 | DJ624 $P_{asr(-250+31)}$-*mSc* (*asr* promoter from -250 to +31 relative to transcription start site inserted upstream of *mSc*) | This study; OK523 + P1 (JM2134b) |

Table S2. Plasmids used in this study

| Plasmid name | Relevant characteristics | Source |
|---|---|---|
| pHDB3 | $Amp^R$, pBR322 derivative, vector control for the genomic DNA library | (6) |
| pRstAB (lab name: pJ2) | $Amp^R$, pHDB3 derivative, carries 'folM-ydgC-rstAB-tus' | This study |
| $pRstAB_{Flag}$ | $Amp^R$, pRstAB derivative. Insertion of 3×FLAG sequence before *rstA* stop codon. | This study |
| pRstABmut1 | $Amp^R$, pRstAB derivative. Directed mutagenesis to substitute $26^{th}$ and $27^{th}$ codons of *rstA* with stop codons. | This study |
| pRstABmut2 | $Amp^R$, pRstAB derivative. Directed mutagenesis to substitute $42^{th}$ and $43^{th}$ codons of *rstB* with stop codons. | This study |
| pRstABmut3 | $Amp^R$, pRstAB derivative. Directed mutagenesis to substitute $19^{th}$ and $20^{th}$ codons of *ydgC* with stop codons. | This study |
| $pRstAB^{LA}$ | $Amp^R$, pRstAB derivative. Site-directed mutagenesis to introduce Y232L and R233A substitutions in *rstB*. | This study |
| pOmpR | $Amp^R$, pHDB3 derivative, carries *'envZ-ompR-greB-yhgF'* region (from positions 3,534,972-3,537,599 on *E. coli* NC_000913.3 genome) | (2) |
| pUT18C | $Amp^R$, plasmid for 2-hybrid assay, carries the T18 cyclase domain | N. Dautin |
| pKT25 | $Kan^R$, plasmid for 2-hybrid assay, carries the T25 cyclase domain | N. Dautin |
| pUT18C-zip | $Amp^R$, pUT18C derivative, plasmid for 2-hybrid assay, carries the T18-Zip positive control | N. Dautin |
| pKT25-zip | $Kan^R$, pKT25 derivative, plasmid for 2-hybrid assay, carries the T25-Zip positive control | N. Dautin |
| pKT25-RstB | $Kan^R$ pKT25 derivative, $P_{lac}$-*cyaA(T25)-rstB* | (7) |
| pKT25-EnvZ | $Kan^R$ pKT25 derivative, $P_{lac}$-*cyaA(T25)-envZ* | (7) |
| pKT25-PhoQ | $Kan^R$ pKT25 derivative, $P_{lac}$-*cyaA(T25)-phoQ* | (8) |
| pUT18C-MgrB | $Amp^R$, pUT18C derivative, $P_{lac}$-*cyaA(T18)-mgrB* | (8) |
| pUT18C-YdgU | $Amp^R$, pUT18C derivative, $P_{lac}$-*cyaA(T18)-ydgU* (cloned using BamHI and XbaI) | This study |
| pUT18C-Asr | $Amp^R$, pUT18C derivative, $P_{lac}$-*cyaA(T18)-asr* (cloned using BamHI and XbaI) without its addressing peptide | This study |
| pUT18C-OmpR | $Amp^R$, pUT18C derivative, $P_{lac}$-*cyaA(T18)-ompR* (cloned using BamHI and XbaI) | This study |
| pUT18C-RstA | $Amp^R$, pUT18C derivative, $P_{lac}$-*cyaA(T18)-rstA* (cloned using BamHI and XbaI) | This study |
| pRLG770 | ColEI, AmpR, used for cloning omrA, omrB or ompC promoter in front of 5S RNA gene | (9) |
| pRLG-omrB | $Amp^R$, pRLG770 derivative, P*omrB* (-104+50 from transcriptional start site cloned with EcoRI and HindIII) | This study |
| pRLG-asr | $Amp^R$, pRLG770 derivative, P*asr* (-250+50 from transcriptional start site cloned with EcoRI and HindIII) | This study |
| pET28a | $Amp^R$ | Novagen |
| pET28a-RstA | $Amp^R$, pET28a derivative, *rstA* ORF cloned with NdeI and HindIII | This study |
| pOmpR-His | $Kan^R$, OmpR-His overproducing plasmid for protein purification | (10) |

Table S3. Oligonucleotides used in this study.

| Name | 5'→3' sequence | Comment |
|---|---|---|
| **Strain** | | |
| 40nt-rrnBt2-F | GAAGGCGAAGCGGCATGCATTTACGTTGACACCATC GAATGGCGCAGAAGGCCATCCTGA | To increase homology of lacZ fusion (forward) |
| lacZ4-67rev | TAACGCCAGGGTTTTCCCAGTCACGACGTTGTAAAA CGACGGCCAGTGAATCCGTAATCATGGT | To increase homology of lacZ fusion (reverse) |
| AK420 | AACGCATAAATTCTTTTATTACTGCTTCGCCTTTGCT CAC | To increase homology of mSc fusion (reverse) |
| oJM11 (lacZ-17-POmrA) | CAGTGAATCCGTAATCATGGTCATAGCTGTTTCCTGT GTGACCTCTGGGATCTTGATTGTGG | P*omrA-lacZ* rev (JM2111, JM2130, JM2136) |
| oJM10 (rrnBt2-POmrA) | GGCGCAGAAGGCCATCCTGACGGATGGCCTTTTTGC GTTTAATCTGTAACAGTAACCGACAATTTAC | P*omrA-lacZ* for (JM2111) |
| oJM21 (PomrABoxB2_2) | TTAATCTGTAACAGTAACCGACAATAACCTTTGGTTA CACTTTGCCCCTTCATTCCTTTGCGTTTTCTCG | P*omrAmut1-lacZ* for PCR1 (JM2130) |
| oJM20 (PomrABoxB2_1) | GGCGCAGAAGGCCATCCTGACGGATGGCCTTTTTGC GTTTAATCTGTAACAGTAACCG | P*omrAmut1-lacZ* for PCR2 (JM2130)<br>P*omrAmut2-lacZ* for PCR3 (JM2136) |
| oJM35 (PomrAmut2_2) | GGTTACACTTTGCGAAACGCTGTTGCGATTGCTCGCT GGCGAAGAGTC | P*omrAmut2-lacZ* for PCR 1 (JM2136) |
| oJM34 (PomrAmut2_1) | GTTTAATCTGTAACAGTAACCGACAATTTAAACCTTT GGTTACACTTTGCGAAACGC | P*omrAmut2-lacZ* for PCR 2 (JM2136) |
| oJM8 (rrnBt2-POmrB) | GGCGCAGAAGGCCATCCTGACGGATGGCCTTTTTGC GTTTAAGTAAGACAAAAAAAGAGATTGCAAAC | P*omrB-lacZ* (JM2110, JM2132) |
| oJM9 (lacZ-17-POmrB) | CAGTGAATCCGTAATCATGGTCATAGCTGTTTCCTGT GTGACCTCTGGGATCACCACTTTAGC | |
| oJM22 (PomrBBoxB2mut2nts) | GTAAGACAAAAAAAGAGATTGCAAACCCTTGGTTGC ACTTTGCGAAACGCTGTTGCG | P*omrBmut-lacZ* PCR1 (JM2132) |
| oJM14 (mScarletPomrArev) | ACTGCTTCGCCTTTGCTCACCATAGCTGTTTCCTGTG TGACCTCTGGGATCTTGATTG | P*omrA-mSc* (JM2203) |
| oJM15 (mScarletPomrAfor) | CGTTTTGGTCCAACTAGTCCGATGCGACACACATGA CGCGAATCTGTAACAGTAACCGAC | |
| oJM12 (mScarletPomrBrev) | ACTGCTTCGCCTTTGCTCACCATAGCTGTTTCCTGTG TGACCTCTGGGATCACCACTTTAGC | P*omrB-mSc* (JM2113) |
| oJM13 (mScarletPomrBfor) | CGTTTTGGTCCAACTAGTCCGATGCGACACACATGA CGCGAAGTAAGACAAAAAAAGAGATTGC | |
| oJM25 (mScarlet_Asr_for) | CGTTTTGGTCCAACTAGTCCGATGCGACACACATGA CGCGttataagtacccaaataaacggattc | Amplification of P*asr* (JM2134, JM327, JM329, JM331) |
| oJM26 (mScarlet_Asr_rev) | ACTGCTTCGCCTTTGCTCACCATAGCTGTTTCCTGTG TGAgtttttcatttaaccccactgc | Construction of P*asr-mSc*, used with oJM25 (JM2134) |
| oJM74 (drstAB::ccdB-catF) | CGCGGTGTATTGTGACGTTTTTATATCTACCGTGAAT GTTTCACTGGCCGTCGTTTTAC | Δ*rstAB::ccdB-cat* (JM109) |
| oJM75 (drstAB::ccdB-catR) | ATAGCACAGTCGTGGTGACTTGACCATCGTGCGCGT AGTGTTTCGGTCATTGGCATGTTC | |

| | | |
|---|---|---|
| oJM55 (RstABfor) | CGCGGTGTATTGTGACGTTTTATATCTACCGTGAATGTTATGAACACTATCGTATTTGTGG | Construction of *rstAD52A, rstA D52E, rstB**, rstB*$^{LA}$ amplified from pRstAB with corresponding modification (JM112, JM113, JM115, JM175, JM177) |
| oJM56 (RstABrev) | ATAGCACAGTCGTGGTGACTTGACCATCGTGCGCGTAGTGTCAGGCAGAGGTAAATTGC | |
| oJM156 (tetPlacrstAfor) | ATTGATAGTAAGTAAAAACAGCGCGGTGTATTGTGACGTTCCTAATTTTTGTTGACACTCTATC | Amplification tet-P*lac* from MG2000 (4) (JM266) |
| oJM157 (tetPlacrstArev) | AATACGATAGTGTTCATAACATTCACGGTAGATATAAAAGTGCTCAGTATCTTGTTATCC | |
| oJM206 (bleo-ydgU-rev) | TTATTTATACAGTTCGTCCATGCCGCCAGTAGAATGACGGAGTAAAATCAGATAACTACAATCAGG | Construction of *kgtP*-P*asr-asr-ydgU-yfiM*, used with oJM25 (JM327) |
| oJM207 (bleo-asr-rev) | TTATTTATACAGTTCGTCCATGCCGCCAGTAGAATGACGGGCCAGCATTACTGTTGAAAAC | Construction of *kgtP*-P*asr-asr-yfiM*, used with oJM25 (JM329) |
| oJM208 (+36asr-rev) | CAGCAGGTTTTGCCGGTTGAGCAGCGGCAACAACCAG | Construction of *kgtP*-P*asr*-Δ*asr-ydgU-yfiM*, used with oJM25 (JM331) |
| oJM209 (asr+273for) | CTGGTTGTTGCCGCTGCTCAACCGGCAAAACCTGCTG | Construction of *kgtP*-P*asr*-Δ*asr-ydgU-yfiM*, used with oJM206 (JM331) |
| oJM186 (Δasr::ccdB-catF) | GCAGTGGGGTTAAATGAAAAAACAAATTGAGGGTATGACATCACTGGCCGTCGTTTTAC | Δ*asr::ccdB-cat* (JM312) |
| oJM187 (Δasr::ccdB-catR) | TTCAGGCGCGAGGGGGCGCGCCAGCATTACTGTTGAAAACTTTCGGTCATTGGCATGTTC | |
| oJM216 (Δasr-cat-for) | GTTAACCCGATCAAGACTACTATTATTGGTAGCTAAATTTCCAGGCATCAAATAAAACGAAAG | ΔP*asr-asr-ydgU::ccdB-cat* (JM335) |
| oJM217 (ΔydgU-cat-rev) | GGGACGGGATGAAAGTGGGAGTAAAATCAGATAACTACAACTTAAAAAAATTACGCCCCGC | |
| **Plasmids** | | |
| oJM44 (pJ2_RstA-flag_for) | TGATATCGACTACAAAGATGACGACGATAAATAGTAAGCGATGAAAAAACTGTTTATCC | pRstAB$_{flag}$ |
| oJM45 (pJ2_RstA-flag_rev) | TGATCTTTATAATCACCGTCATGGTCTTTGTAGTCTTCCCATGCATGAGGCG | |
| oJM1 (KOrstAfor) | GCGTACCTGGCAAAACATGAT**TAAT**AGGTTACCGTAGAGCCGCGC | pRstABmut1 |
| oJM2 (KOrstArev) | GCGCGGCTCTACGGTAACCT**ATTA**ATCATGTTTTGCCAGGTACGC | |
| oJM64 (pJ2mut2_for) | GGGCAAACAGTCGCTGGATGATT**AATA**GAACAGTTCGCTGTATCTGATGC | pRstABmut2 |
| oJM65 (pJ2mut2_rev) | GCATCAGATACAGCGAACTGTTC**TATT**AATCATCCAGCGACTGTTTGCCC | |
| oJM170 (pJ2mut3for) | CGGCGATATAATAATTTTTCGTTTTTGCT**TATTA**ACCAATCAACAGCACTACCAGC | pRstABmut3 |

| | | |
|---|---|---|
| oJM171 (pJ2mut3rev) | GCTGGTAGTGCTGTTGATTGGT**TAATA**AGCAAAAACGAAAAATTATTATATCGCCG | |
| oJM122 (pJ2rstBLAfor) | CCGTTAGTGCGCCTGCGT**CTGGC**ACTGGAGATGAGCGATAACC | pRstAB$^{LA}$ |
| oJM123 (pJ2rstBLArev) | GGTTATCGCTCATCTCCAGT**GCCAG**ACGCAGGCGCACTAACGG | |
| oJM28 (RstAD52Afor) | GGATTTGGTGTTACTCG**CG**ATCATGCTACCAGGCAAGGAC | pRstAB_D52A |
| oJM29 (RstAD52Arev) | GTCCTTGCCTGGTAGCATGAT**CG**CGAGTAACACCAAATCC | |
| oJM30 (RstAD52Efor) | GGATTTGGTGTTACTCGA**A**ATCATGCTACCAGGCAAGGAC | pRstAB_D52E |
| oJM31 (RstAD52Erev) | GTCCTTGCCTGGTAGCATGAT**T**TCGAGTAACACCAAATCC | |
| oJM198 (XbaI-ydgU-for) | CGTCTAGACATGGTGGGCCGTTATCG | pUT18C-YdgU |
| oJM199 (BamHI-ydgU-rev) | GCGGATCCTCAGGAAAGATAAAAACGGGC | |
| oJM196 (XbaI-asr-for) | CGTCTAGACGCAGAGACTACGACCAC | pUT18C-Asr |
| oJM197 (BamHI-asr-rev) | GCGGATCCTTACGCTGCGGGTTGTG | |
| oJM66 (XbaIrstAfor) | GACTCTAGACATGAACACTATCGTATTTGTGGAAG | pUT18C-RstA |
| oJM67 (BamHIrstArev) | GTCGGATCCGCTTCCCATGCATGAGGCGC | |
| XbaI-OmpR_for | GCATTCTAGACATGCAAGAGAACTACAAGATTCTGG | pUT18C-OmpR |
| oJM68 (BamHIompRrev) | GCATGGATCCGCTTTAGAGCCGTCCGG | |
| oJM23 (NdeIRstAfor) | CGAGCATATGaacactatcgtatttgtgg | pET28a-RstA |
| oJM24 (HindIIIRstArev) | CGACAAGCTTacagtttttcatcgcTTAttcc | |
| oJM54 (Eco-104OmrB) | GCAGAATTCAAGTAAGACAAAAAAAGAGATTGCAAACC | pRLG-omrB |
| OmrB+50Hind | CGAAAGCTTTAATTCATGTGCTCAACCCGAAG | |
| oJM51 (asr+50-Hind) | CGAAAGCTTATTGTCATACCCTCAATTTGTTTTTTC | pRLG-asr |
| oJM52 (Eco-250asr) | GCAGAATTCTTATAAGTACCCAAATAAACGGATTC | |
| oJM66 (XbaIrstAfor) | GACTCTAGACATGAACACTATCGTATTTGTGGAAG | pUT18C-RstA |
| oJM67 (BamHIrstArev) | GCATGGATCCGCTTTAGAGCCGTCCGG | |
| XbaI-OmpR_for | GCATTCTAGACATGCAAGAGAACTACAAGATTCTGG | pUT18C-OmpR |

| | | |
|---|---|---|
| oJM68 (BamHIompRrev) | GCATGGATCCGCTTTAGAGCCGTCCGG | |
| oJM196 (XbaI-asr-for) | CGTCTAGACGCAGAGACTACGACCAC | pUT18C-Asr |
| oJM197 (BamHI-asr-rev) | CGTCTAGACGCAGAGACTACGACCAC | |
| oJM198 (XbaI-ydgU-for) | GCGGATCCTTACGCTGCGGGTTGTG | pUT18C-ydgU |
| oJM199 (BamHI-ydgU-rev) | GCGGATCCTCAGGAAAGATAAAAACGGGC | |
| **DNA used in DRaCALA** | | |
| oJM89 (pRLG770MCSfor) | CCTTTCGTCTTCAAGAATTC | Primer forward commun |
| oJM90 (omrB+8-8) | CCTCTGGGATCACCAC | Primer rev for P*omrB* |
| oJM51 (asr+50-Hind) | CGAAAGCTTATTGTCATACCCTCAATTTGTTTTTC | Primer rev for P*asr* |
| **Northern Blot probe** | | |
| OmrAmut-probe | (Bio)CAGGTTGGTGCAAGAGACAGGGTACGAAGAGCGTACCG | |
| OmrBmut-probe | (Bio)CGCAGGCTGGTGTAATTCATGTGCTCAACCCGAAGTTGA | |
| ssrA-probe | (Bio)CGCCACTAACAAACTAGCCTGATTAAGTTTTAACGCTTCA | |
| asr2-probe | (Bio)CAGAAGACAGACCCATAGCAGCGGCAACAACCAGAGCTA | |

Table S4. *Enterobacteriaceae* used in Fig. 1B

| Name | NCBI_accession_number |
|---|---|
| Escherichia coli | NC_000913.3 |
| Escherichia coli | NC_011993.1 |
| Citrobacter freundii | NZ_CP033744.1 |
| Salmonella enterica | NC_003197.2 |
| Shigella dysenteriae | NZ_CP061527.1 |
| Klebsiella aerogenes | NZ_LR134475.1 |
| Klebsiella pneumoniae | CP003200.1 |
| Enterobacter hormaechei | CP041054.1 |
| Enterobacter kobei | CP017181.1 |
| Enterobacter ludwigii | CP017279.1 |
| Enterobacter cancerogenus | CP025225.1 |
| Shigella sonnei | NZ_NQBD01000044.1 |
| Cronobacter sakazakii | NZ_CP027107.1 |
| Kosakonia sacchari | CP007215.3 |
| Buttiauxella agrestis | AP023184.1 |
| Shimwellia blattae | CP001560.1 |

| | |
|---|---|
| Metakosakonia sp. | AP018756.1 |
| Lelliottia jeotgali | CP018628.1 |
| Lelliottia amnigena | CP015774.2 |
| Lelliottia nimipressuralis | CP025034.2 |
| Lelliottia sp. | CP028520.1 |
| Cedecea lapagei | LR134201.1 |
| Leclercia sp. | CP026167.1 |
| Raoultella ornithinolytica | CP004142.1 |
| Kluyvera intermedia | LR134138.1 |
| Kluyvera ascorbata | AP022665.1 |
| Yokenella regensburgei | CP050811.1 |
| Buttiauxella sp. | CP033076.1 |
| Cronobacter turicensis | FN543093.2 |
| Cronobacter malonaticus | CP006731.1 |
| Atlantibacter hermannii | LR134136.1 |

Table S5. *Enterobacterales* used in Fig. 6C

| Nom_espece | Numero_acces_GCF | Famille |
|---|---|---|
| Escherichia_coli_K12_MG1655 | GCF_000005845.2 | Enterobacteriaceae |
| Escherichia_coli_LF82 | GCF_000284495.1 | Enterobacteriaceae |
| Escherichia_coli_O157H7_EDL933 | GCF_000006665.1 | Enterobacteriaceae |
| Shigella_flexneri_2a_301 | GCF_000006925.2 | Enterobacteriaceae |
| Shigella_dysenteriae_Sd197 | GCF_000012005.1 | Enterobacteriaceae |
| Shigella_sonnei_ATCC_29930 | GCF_002950395.1 | Enterobacteriaceae |
| Salmonella_enterica_Typhimurium_LT2 | GCF_000006945.2 | Enterobacteriaceae |
| Salmonella_enterica_Typhi_Ty2 | GCF_000007545.1 | Enterobacteriaceae |
| Salmonella_bongori_NCTC_12419 | GCF_000252995.1 | Enterobacteriaceae |
| Citrobacter_amalonaticus_FDAARGOS_1489 | GCF_020099335.1 | Enterobacteriaceae |
| Citrobacter_rodentium_DSM_16636 | GCF_021278985.1 | Enterobacteriaceae |
| Citrobacter_farmeri_FDAARGOS_1423 | GCF_019048065.1 | Enterobacteriaceae |
| Citrobacter_pasteurii_FDAARGOS_1424 | GCF_019047765.1 | Enterobacteriaceae |
| Citrobacter_arsenatis_LY-1 | GCF_004353845.1 | Enterobacteriaceae |
| Citrobacter_tructae_SNU_WT2 | GCF_004684345.1 | Enterobacteriaceae |
| Citrobacter_freundii_MSB1_1H | GCF_904859905.1 | Enterobacteriaceae |
| Citrobacter_portucalensis_Cf7303 | GCF_023374935.1 | Enterobacteriaceae |
| Citrobacter_braakii_MiY-A | GCF_009648935.1 | Enterobacteriaceae |
| Citrobacter_koseri_ATCC_BAA-895 | GCF_000018045.1 | Enterobacteriaceae |
| Citrobacter_werkmanii_FDAARGOS_616 | GCF_008693645.1 | Enterobacteriaceae |
| Citrobacter_sedlakii_3347689II | GCF_018128425.1 | Enterobacteriaceae |
| Klebsiella_pneumoniae_MGH78578 | GCF_000016305.1 | Enterobacteriaceae |
| Klebsiella_oxytoca_NCTC13727 | GCF_900636985.1 | Enterobacteriaceae |
| Klebsiella_quasipneumoniae_FDAARGOS_1503 | GCF_020099175.1 | Enterobacteriaceae |

| | | |
|---|---|---|
| Klebsiella_variicola_F2R9T | GCF_020525545.1 | Enterobacteriaceae |
| Klebsiella_electrica_DSM_102253 | GCF_006711645.1 | Enterobacteriaceae |
| Klebsiella_huaxiensis_WCHKl090001 | GCF_003261575.2 | Enterobacteriaceae |
| Klebsiella_africana_200023 | GCF_020526085.1 | Enterobacteriaceae |
| Klebsiella_pneumoniae_HS11286 | GCF_000240185.1 | Enterobacteriaceae |
| Klebsiella_michiganensis_THO-011 | GCF_015139575.1 | Enterobacteriaceae |
| Klebsiella_grimontii_2750 | GCF_042137965.1 | Enterobacteriaceae |
| Klebsiella_pasteurii_Sb-24 | GCF_018139045.1 | Enterobacteriaceae |
| Klebsiella_quasivariicola_08A119 | GCF_020525665.1 | Enterobacteriaceae |
| Klebsiella_spallanzanii | GCF_902158555.1 | Enterobacteriaceae |
| Klebsiella_indica_TOUT106 | GCF_005860775.1 | Enterobacteriaceae |
| Raoultella_ornithinolytica_RoM27LC23 | GCF_030505655.1 | Enterobacteriaceae |
| Raoultella_planticola_FDAARGOS_64 | GCF_000783935.2 | Enterobacteriaceae |
| Raoultella_terrigena_JH01 | GCF_012029655.1 | Enterobacteriaceae |
| Raoultella_lignicola_TW_WC1a.1 | GCF_036561965.2 | Enterobacteriaceae |
| Raoultella_scottii_Txe2.2 | GCF_036561945.1 | Enterobacteriaceae |
| Enterobacter_cloacae_ATCC_13047 | GCF_000025565.1 | Enterobacteriaceae |
| Enterobacter_hormaechei_FDAARGOS_1433 | GCF_019048245.1 | Enterobacteriaceae |
| Enterobacter_roggenkampii_DSM_16690 | GCF_001729805.1 | Enterobacteriaceae |
| Enterobacter_ludwigii_EN-119 | GCF_001750725.1 | Enterobacteriaceae |
| Enterobacter_bugandensis_EB-247 | GCF_900324475.1 | Enterobacteriaceae |
| Enterobacter_oligotrophicus_CCA6 | GCF_009176645.1 | Enterobacteriaceae |
| Enterobacter_chengduensis_WCHECl-C4 | GCF_001984825.2 | Enterobacteriaceae |
| Enterobacter_huaxiensis_090008 | GCF_003594935.2 | Enterobacteriaceae |
| Enterobacter_adelaidei_ECC3473 | GCF_041937165.1 | Enterobacteriaceae |
| Enterobacter_dykesii_E1 | GCF_008364625.2 | Enterobacteriaceae |
| Enterobacter_cloacae_1382 | GCF_905331265.2 | Enterobacteriaceae |
| Enterobacter_asburiae_17Nkhm-UP2 | GCF_007035805.1 | Enterobacteriaceae |
| Enterobacter_kobei_11778-yvys | GCF_023023125.1 | Enterobacteriaceae |
| Enterobacter_mori_ACYC.E9L | GCF_022014715.1 | Enterobacteriaceae |
| Enterobacter_soli | GCF_000224675.1 | Enterobacteriaceae |
| Kluyvera_intermedia_N2-1 | GCF_009649915.1 | Enterobacteriaceae |
| Kluyvera_ascorbata_SK | GCF_023195735.1 | Enterobacteriaceae |
| Kluyvera_sichuanensis_lhn-g4 | GCF_040930405.1 | Enterobacteriaceae |
| Kluyvera_cryocrescens_NBRC_102467 | GCF_001571285.1 | Enterobacteriaceae |
| Kluyvera_georgiana_WCH1410 | GCF_001682915.1 | Enterobacteriaceae |
| Leclercia_pneumoniae_49125 | GCF_017348915.1 | Enterobacteriaceae |
| Leclercia_adecarboxylata_USDA-ARS-USMARC-60222 | GCF_001518835.1 | Enterobacteriaceae |
| Leclercia_tamurae_H6S3 | GCF_025566055.1 | Enterobacteriaceae |
| Leclercia_barmai_EMC7 | GCF_019913755.1 | Enterobacteriaceae |
| Lelliottia_nimipressuralis_SCAID_67 | GCF_002211585.1 | Enterobacteriaceae |

| | | |
|---|---|---|
| Lelliottia_amnigena_FDAARGOS_424 | GCF_002073755.2 | Enterobacteriaceae |
| Cronobacter_sakazakii_ATCC_BAA-894 | GCF_000017665.1 | Enterobacteriaceae |
| Franconibacter_pulveris_DJ34 | GCF_001077855.1 | Enterobacteriaceae |
| Pluralibacter_gergoviae_FB2 | GCF_003064215.1 | Enterobacteriaceae |
| Siccibacter_turicensis_493 | GCF_004168465.1 | Enterobacteriaceae |
| Cedecea_lapagei_NCTC11466 | GCF_900635955.1 | Enterobacteriaceae |
| Cedecea_davisae_739Q | GCF_044095835.1 | Enterobacteriaceae |
| Cedecea_neteri_FDAARGOS_392 | GCF_002393445.1 | Enterobacteriaceae |
| Cedecea_sulfonylureivorans_LAM2020 | GCF_016756775.1 | Enterobacteriaceae |
| Cedecea_colo_ZA | GCF_011808225.1 | Enterobacteriaceae |
| Kosakonia_sacchari_SP1 | GCF_000300455.3 | Enterobacteriaceae |
| Shimwellia_blattae_DSM_4481 | GCF_000262305.1 | Enterobacteriaceae |
| Buttiauxella_ferragutiae_H4-C11 | GCF_022637515.1 | Enterobacteriaceae |
| Buttiauxella_agrestis_NCTC12119 | GCF_900446255.1 | Enterobacteriaceae |
| Buttiauxella_gaviniae_ATCC_51604 | GCF_001654835.1 | Enterobacteriaceae |
| Buttiauxella_massiliensis_Marseille-P9829 | GCF_902500225.1 | Enterobacteriaceae |
| Buttiauxella_brennerae_ATCC_51605 | GCF_001654925.1 | Enterobacteriaceae |
| Buttiauxella_izardii_CCUG_35510 | GCF_003601925.1 | Enterobacteriaceae |
| Buttiauxella_warmboldiae_CCUG_35512 | GCF_003818135.1 | Enterobacteriaceae |
| Buttiauxella_noackiae_MCE | GCF_000737905.1 | Enterobacteriaceae |
| Trabulsiella_odontotermitis_TBY01 | GCF_030053895.1 | Enterobacteriaceae |
| Trabulsiella_guamensis_ATCC_49490 | GCF_000734965.1 | Enterobacteriaceae |
| Yokenella_regensburgei_DSM_5079 | GCF_003634235.1 | Enterobacteriaceae |
| Erwinia_amylovora_CFBP1430 | GCF_000091565.1 | Erwiniaceae |
| Erwinia_pyrifoliae_DSM_12163 | GCF_000026985.1 | Erwiniaceae |
| Erwinia_pyrifoliae_EpK1-15 | GCF_002952315.1 | Erwiniaceae |
| Erwinia_rhapontici_BY21311 | GCF_020683125.1 | Erwiniaceae |
| Erwinia_tracheiphila_BHKY | GCF_021365465.1 | Erwiniaceae |
| Candidatus_Erwinia_haradaeae_ErCicurvipes | GCF_900698925.1 | Erwiniaceae |
| Erwinia_piriflorinigrans_CFBP_5888 | GCF_001050515.1 | Erwiniaceae |
| Erwinia_phyllosphaerae_CMYE1 | GCF_019132875.1 | Erwiniaceae |
| Erwinia_aeris_ACCC_02193 | GCF_041224955.1 | Erwiniaceae |
| Erwinia_psidii_IBSBF_435 | GCF_003846135.1 | Erwiniaceae |
| Erwinia_oleae_DAPP-PG531 | GCF_000770305.1 | Erwiniaceae |
| Candidatus_Erwinia_dacicola_IL | GCF_001756855.1 | Erwiniaceae |
| Erwinia_aphidicola_USMM130 | GCF_037149315.1 | Erwiniaceae |
| Erwinia_mallotivora_BT-MARDI | GCF_000590885.1 | Erwiniaceae |
| Erwinia_plantamica_OPT-41 | GCF_043420595.1 | Erwiniaceae |
| Pantoea_ananatis_PA13 | GCF_000233595.1 | Erwiniaceae |
| Pantoea_stewartii_ZJ-FGZX1 | GCF_011044475.1 | Erwiniaceae |
| Pantoea_agglomerans_FDAARGOS_1447 | GCF_019048385.1 | Erwiniaceae |
| Pantoea_vagans_LMG_24199 | GCF_004792415.1 | Erwiniaceae |

| | | |
|---|---|---|
| Tatumella_ptyseos_ATCC_33301 | GCF_000439895.1 | Erwiniaceae |
| Phaseolibacter_flectens_ATCC_12775 | GCF_000686145.1 | Erwiniaceae |
| Wigglesworthia_glossinidia_morsitans | GCF_000247565.1 | Erwiniaceae |
| Paramixta_manurensis_PD-1 | GCF_013285385.1 | Erwiniaceae |
| Mixta_gavinia_DSM_22758 | GCF_002953195.1 | Erwiniaceae |
| Mixta_hanseatica_X22927 | GCF_023517775.1 | Erwiniaceae |
| Buchnera_aphidicola_Sg | GCF_000007365.1 | Erwiniaceae |
| Pectobacterium_carotovorum_PC1 | GCF_000023605.1 | Pectobacteriaceae |
| Pectobacterium_atrosepticum_SCRI1043 | GCF_000011605.1 | Pectobacteriaceae |
| Pectobacterium_parmentieri_RNS_08-42-1A | GCF_001742145.1 | Pectobacteriaceae |
| Pectobacterium_punjabense_SS95 | GCF_012427845.1 | Pectobacteriaceae |
| Pectobacterium_wasabiae_CFBP_3304 | GCF_001742185.1 | Pectobacteriaceae |
| Pectobacterium_colocasium_LJ1 | GCF_020181655.1 | Pectobacteriaceae |
| Pectobacterium_cacticida_CFBP3628 | GCF_036885195.1 | Pectobacteriaceae |
| Pectobacterium_araliae_MAFF_302110 | GCF_037076465.1 | Pectobacteriaceae |
| Pectobacterium_quasiaquaticum_A477-S1-J17 | GCF_014946775.2 | Pectobacteriaceae |
| Pectobacterium_aquaticum_A212-S19-A16 | GCF_003382565.3 | Pectobacteriaceae |
| Pectobacterium_carotovorum_WPP14 | GCF_013488025.1 | Pectobacteriaceae |
| Pectobacterium_brasiliense_1692 | GCF_009873295.1 | Pectobacteriaceae |
| Pectobacterium_atrosepticum_21A | GCF_000740965.1 | Pectobacteriaceae |
| Pectobacterium_aroidearum_L6 | GCF_015689195.1 | Pectobacteriaceae |
| Pectobacterium_odoriferum_JK2.1 | GCF_009931295.1 | Pectobacteriaceae |
| Pectobacterium_versatile_14A | GCF_003932035.1 | Pectobacteriaceae |
| Pectobacterium_polaris_NIBIO1392 | GCF_002288545.1 | Pectobacteriaceae |
| Pectobacterium_parvum_FN20211 | GCF_020971565.1 | Pectobacteriaceae |
| Pectobacterium_peruviense_A350-S18-N16 | GCF_003312355.2 | Pectobacteriaceae |
| Pectobacterium_actinidiae_KKH3 | GCF_000803315.1 | Pectobacteriaceae |
| Pectobacterium_polonicum_DPMP315 | GCF_005497185.1 | Pectobacteriaceae |
| Pectobacterium_zantedeschiae_9M | GCF_004137795.1 | Pectobacteriaceae |
| Dickeya_dadantii_3937 | GCF_000147055.1 | Pectobacteriaceae |
| Dickeya_dadantii_DSM_18020 | GCF_003049785.1 | Pectobacteriaceae |
| Dickeya_zeae_MS2 | GCF_002887555.1 | Pectobacteriaceae |
| Dickeya_fangzhongdai_DSM_101947 | GCF_002812485.1 | Pectobacteriaceae |
| Dickeya_solani_IPO_2222 | GCF_001644705.1 | Pectobacteriaceae |
| Dickeya_poaceiphila_NCPPB_569 | GCF_007858975.2 | Pectobacteriaceae |
| Dickeya_aquatica_174-2 | GCF_900095885.1 | Pectobacteriaceae |
| Dickeya_dianthicola_ME23 | GCF_003403135.1 | Pectobacteriaceae |
| Dickeya_chrysanthemi_Ech1591 | GCF_000023565.1 | Pectobacteriaceae |
| Dickeya_oryzae_A003-S1-M15 | GCF_020406815.2 | Pectobacteriaceae |
| Dickeya_parazeae_Ech586 | GCF_000025065.1 | Pectobacteriaceae |
| Dickeya_lacustris_S29 | GCF_003934295.1 | Pectobacteriaceae |
| Dickeya_undicola_2B12 | GCF_000784735.1 | Pectobacteriaceae |

| | | |
|---|---|---|
| Brenneria_goodwinii_FRB141 | GCF_002291445.1 | Pectobacteriaceae |
| Brenneria_izadpanahii_Iran_50 | GCF_017569925.1 | Pectobacteriaceae |
| Brenneria_uluponensis_K61 | GCF_032395385.1 | Pectobacteriaceae |
| Brenneria_nigrifluens_DSM_30175 | GCF_005484965.1 | Pectobacteriaceae |
| Prodigiosinella_confusarubida_ATCC_39006 | GCF_000463345.2 | Pectobacteriaceae |
| Lonsdalea_populi_N-5-1 | GCF_015999465.1 | Pectobacteriaceae |
| Lonsdalea_britannica_477 | GCF_003515985.1 | Pectobacteriaceae |
| Sodalis_glossinidius_morsitans | GCF_000010085.1 | Pectobacteriaceae |
| Candidatus_Sodalis_pierantonius_SOPE | GCF_000517405.1 | Pectobacteriaceae |
| Sodalis_ligni_159R | GCF_004346745.1 | Pectobacteriaceae |
| Candidatus_Sodalis_endolongispinus_SOD | GCF_018777395.1 | Pectobacteriaceae |
| Sodalis_like_endosymbiont_SPI1 | GCF_001602625.1 | Pectobacteriaceae |
| Musicola_paradisiaca_Ech703 | GCF_000023545.1 | Pectobacteriaceae |
| Musicola_keenii_A3967 | GCF_014855505.1 | Pectobacteriaceae |
| Yersinia_pestis_CO92 | GCF_000009065.1 | Yersiniaceae |
| Yersinia_entomophaga_MH96 | GCF_001656035.1 | Yersiniaceae |
| Yersinia_enterocolitica_Y11 | GCF_000253175.1 | Yersiniaceae |
| Yersinia_pseudotuberculosis_NCTC10275 | GCA_900637475.1 | Yersiniaceae |
| Yersinia_similis_Y_sim_228 | GCF_000582515.1 | Yersiniaceae |
| Yersinia_mollaretii_ATCC_43969 | GCF_013282725.1 | Yersiniaceae |
| Yersinia_hibernica_CFS1934 | GCF_004124235.1 | Yersiniaceae |
| Yersinia_canaria_NCTC_14382 | GCF_009831415.1 | Yersiniaceae |
| Yersinia_pestis_A1122 | GCF_000222975.1 | Yersiniaceae |
| Yersinia_ruckeri_KMM821 | GCF_017498685.1 | Yersiniaceae |
| Yersinia_intermedia_FDAARGOS_730 | GCF_009730055.1 | Yersiniaceae |
| Yersinia_rohdei_YRA | GCF_000834455.1 | Yersiniaceae |
| Yersinia_aldovae_670-83 | GCF_000834395.1 | Yersiniaceae |
| Yersinia_bercovieri_Y195 | GCF_037060685.1 | Yersiniaceae |
| Yersinia_alsatica_SCPM-O-B-7604 | GCF_025133195.1 | Yersiniaceae |
| Yersinia_vastinensis_SCPM-O-B-3977 | GCF_044905075.1 | Yersiniaceae |
| Yersinia_rochesterensis_ATCC_BAA-2637 | GCF_003600645.1 | Yersiniaceae |
| Yersinia_kristensenii_NCTC11471 | GCA_900460525.1 | Yersiniaceae |
| Yersinia_frederiksenii_ATCC_33641 | GCA_000754805.1 | Yersiniaceae |
| Yersinia_massiliensis_CCUG_53443 | GCF_000312485.1 | Yersiniaceae |
| Yersinia_proxima_IP37424 | GCF_902170785.1 | Yersiniaceae |
| Serratia_fonticola_DSM_4576 | GCF_001006005.1 | Yersiniaceae |
| Serratia_liquefaciens_ATCC_27592 | GCF_000422085.1 | Yersiniaceae |
| Serratia_rubidaea_FDAARGOS_926 | GCF_016026735.1 | Yersiniaceae |
| Serratia_symbiotica_CWBI-2.3 | GCF_000821185.2 | Yersiniaceae |
| Serratia_entomophila_A1 | GCF_021462285.1 | Yersiniaceae |
| Serratia_nevei_LMG_31536 | GCF_037948395.1 | Yersiniaceae |
| Serratia_ficaria_NCTC12148 | GCA_900187015.1 | Yersiniaceae |

| | | |
|---|---|---|
| Serratia_surfactantfaciens_YD25 | GCF_001642805.2 | Yersiniaceae |
| Serratia_aquatilis_2015-2462-01 | GCF_044865145.1 | Yersiniaceae |
| Serratia_rhizosphaerae_KUDC3025 | GCF_009817885.1 | Yersiniaceae |
| Serratia_marcescens_ELP1.10 | GCF_030291735.1 | Yersiniaceae |
| Serratia_plymuthica_AS9 | GCF_000214235.1 | Yersiniaceae |
| Serratia_ureilytica_T6 | GCF_017309605.1 | Yersiniaceae |
| Serratia_nematodiphila_DH-S01 | GCF_004768745.1 | Yersiniaceae |
| Serratia_proteamaculans_EBP3064 | GCF_949794035.1 | Yersiniaceae |
| Serratia_quinivorans_NCTC13188 | GCA_900638135.1 | Yersiniaceae |
| Serratia_sarumanii_K-M0228 | GCF_035749905.1 | Yersiniaceae |
| Serratia_grimesii_BXF1 | GCF_900186025.1 | Yersiniaceae |
| Serratia_inhibens_S40 | GCF_003591175.1 | Yersiniaceae |
| Serratia_oryzae_J11-6 | GCF_001976145.1 | Yersiniaceae |
| Chania_multitudinisentens_RB-25 | GCF_000520015.2 | Yersiniaceae |
| Rahnella_aquatilis_HX2 | GCF_000255535.1 | Yersiniaceae |
| Rouxiella_badensis_DAR84756 | GCF_026967515.2 | Yersiniaceae |
| Rouxiella_chamberiensis_130333 | GCF_000951135.1 | Yersiniaceae |
| Rouxiella_aceri_SAP-1 | GCF_012933545.1 | Yersiniaceae |
| Rouxiella_silvae_213 | GCF_002093625.1 | Yersiniaceae |
| Ewingella_americana_CCUG_14506T | GCF_008693045.1 | Yersiniaceae |
| Hafnia_alvei_A23BA | GCF_011617105.1 | Hafniaceae |
| Hafnia_paralvei_AVS0177 | GCF_020150375.1 | Hafniaceae |
| Hafnia_psychrotolerans_CGMCC_1.12806 | GCF_014639435.1 | Hafniaceae |
| Edwardsiella_tarda_FDAARGOS_1473 | GCF_019933175.1 | Hafniaceae |
| Edwardsiella_anguillarum_080813 | GCF_000264765.2 | Hafniaceae |
| Edwardsiella_hoshinae_FDAARGOS_940 | GCF_016026395.1 | Hafniaceae |
| Edwardsiella_piscicida_18EpOKYJ | GCF_021733145.1 | Hafniaceae |
| Edwardsiella_ictaluri_S07-698 | GCF_003074995.2 | Hafniaceae |
| Enterobacillus_tribolii_DSM_103736 | GCF_003363015.1 | Hafniaceae |
| Obesumbacterium_proteus_LE8 | GCF_000980985.1 | Hafniaceae |
| Morganella_morganii_ATCC_25830 | GCF_006094455.1 | Morganellaceae |
| Proteus_myxofaciens_ATCC_19692 | GCF_001654855.1 | Morganellaceae |
| Proteus_appendicitis_HZ0627 | GCF_030271835.1 | Morganellaceae |
| Proteus_mirabilis_HI4320 | GCF_000069965.1 | Morganellaceae |
| Proteus_vulgaris_USDA-ARS-USMARC-49741 | GCF_025200655.1 | Morganellaceae |
| Proteus_terrae_ZN2 | GCF_011045835.1 | Morganellaceae |
| Proteus_penneri_S178-2 | GCF_022369495.1 | Morganellaceae |
| Providencia_stuartii_MRSN_2154 | GCF_000259175.1 | Morganellaceae |
| Providencia_rettgeri_FDAARGOS_1450 | GCF_019048105.1 | Morganellaceae |
| Providencia_heimbachae_NCTC12003 | GCA_900475855.1 | Morganellaceae |
| Providencia_manganoxydans_LLDRA6 | GCF_016618195.1 | Morganellaceae |
| Providencia_hangzhouensis_PR-310 | GCF_029193595.2 | Morganellaceae |

| | | |
|---|---|---|
| Providencia_zhijiangensis_D4759 | GCF_030315915.2 | Morganellaceae |
| Providencia_stuartii_41 | GCF_035747985.1 | Morganellaceae |
| Photorhabdus_laumondii_TT01 | GCF_003343245.1 | Morganellaceae |
| Photorhabdus_thracensis_DSM_15199 | GCF_001010285.1 | Morganellaceae |
| Photorhabdus_asymbiotica_ATCC43949 | GCF_000196475.1 | Morganellaceae |
| Arsenophonus_nasoniae_FIN | GCF_004768525.1 | Morganellaceae |
| Arsenophonus_apicola_ArsBeeUS | GCF_020268605.1 | Morganellaceae |
| Xenorhabdus_nematophila_ATCC_19061 | GCF_000252955.1 | Morganellaceae |
| Xenorhabdus_doucetiae_FRM16 | GCF_000968195.1 | Morganellaceae |
| Xenorhabdus_griffiniae_Kalro | GCF_030016155.2 | Morganellaceae |
| Moellerella_wisconsensis_ATCC_35017 | GCF_000768415.1 | Morganellaceae |
| Budvicia_aquatica_FDAARGOS_387 | GCF_002591785.1 | Budviciaceae |
| Budvicia_diplopodorum_D9 | GCF_009800925.1 | Budviciaceae |
| Leminorella_grimontii_LG-KP-E1-2-T0 | GCF_039789245.1 | Budviciaceae |
| Leminorella_richardii_NCTC12151 | GCA_900478135.1 | Budviciaceae |
| Pragia_fontium_NCTC12284 | GCA_900638655.1 | Budviciaceae |
| Limnobaculum_zhutongyuii_CF-458 | GCF_004295645.1 | Budviciaceae |
| Limnobaculum_parvum_HYN0051 | GCF_003096015.2 | Budviciaceae |
| Limnobaculum_xujianqingii_CF-1111 | GCF_013394855.1 | Budviciaceae |
| Limnobaculum_allomyrinae_BWR-B9 | GCF_016649425.1 | Budviciaceae |
| Limnobaculum_eriocheiris_LJY008 | GCF_021010575.1 | Budviciaceae |
| Pasteurella_multocida_Pm70 | GCF_000006825.1 | Outgroup |
| Haemophilus_influenzae_FDAARGOS_1560 | GCF_020736045.1 | Outgroup |
| Haemophilus_parainfluenzae_FDAARGOS_1000 | GCF_016127215.1 | Outgroup |
| Haemophilus_parahaemolyticus_FDAARGOS_1199 | GCF_016889385.1 | Outgroup |
| Haemophilus_pittmaniae_NCTC13334 | GCA_900186995.1 | Outgroup |
| Haemophilus_aegyptius_NCTC8502 | GCA_900475885.1 | Outgroup |
| Haemophilus_ducreyi_VAN2 | GCF_001647695.1 | Outgroup |
| Actinobacillus_pleuropneumoniae_L20 | GCF_000015885.1 | Outgroup |
| Mannheimia_haemolytica_USDA-ARS-USMARC-191 | GCF_002285575.1 | Outgroup |
| Haemophilus_haemolyticus_NCTC10839 | GCA_900477945.1 | Outgroup |
| Haemophilus_seminalis_SZY_H1 | GCF_006384255.1 | Outgroup |
| Haemophilus_paracuniculus_CCUG_43573 | GCF_002015115.1 | Outgroup |
| Haemophilus_sputorum_HK_2154 | GCF_000287615.1 | Outgroup |
| Vibrio_cholerae_N16961 | GCF_000006745.1 | Outgroup |
| Vibrio_parahaemolyticus_RIMD_2210633 | GCF_000196095.1 | Outgroup |
| Vibrio_vulnificus_CMCP6 | GCF_000039765.1 | Outgroup |
| Vibrio_fischeri_ES114 | GCF_000011805.1 | Outgroup |
| Francisella_halioticida_DSM_23729 | GCF_002211785.1 | Outgroup |
| Francisella_adeliensis_FDC440 | GCF_003290445.1 | Outgroup |
| Francisella_persica_FSC845 | GCF_001275365.1 | Outgroup |

| | | |
|---|---|---|
| Francisella_marina_E95-16 | GCF_008369785.1 | Outgroup |
| Francisella_uliginis_TX07-7310 | GCF_001895265.1 | Outgroup |
| Pseudomonas_putida_NBRC_14164 | GCF_000412675.1 | Outgroup |
| Pseudomonas_chlororaphis_ATCC_9446 | GCF_036689615.1 | Outgroup |
| Pseudomonas_protegens_CHA0 | GCF_900560965.1 | Outgroup |
| Pseudomonas_syringae_DC3000 | GCF_000007805.1 | Outgroup |
| Pseudomonas_entomophila_L48 | GCF_000026105.1 | Outgroup |
| Pseudomonas_cichorii_DSM_50259 | GCF_018343775.1 | Outgroup |
| Pseudomonas_tohonis_TUM18999 | GCF_012767755.2 | Outgroup |
| Pseudomonas_silesiensis_A3 | GCF_001661075.1 | Outgroup |
| Pseudomonas_monsensis_PGSB_8459 | GCF_014268495.2 | Outgroup |
| Pseudomonas_glycineae_MS586 | GCF_001594225.2 | Outgroup |
| Pseudomonas_kribbensis_46-2 | GCF_003352185.1 | Outgroup |
| Pseudomonas_nunensis_In5 | GCF_024296925.1 | Outgroup |
| Pseudomonas_viciae_11K1 | GCF_004786035.1 | Outgroup |
| Pseudomonas_ogarae_SWRI108 | GCF_014268695.2 | Outgroup |
| Pseudomonas_shahriarea_SWRI52 | GCF_014268455.2 | Outgroup |
| Pseudomonas_wenzhouensis_A20 | GCF_021029445.1 | Outgroup |
| Pseudomonas_mucidolens_NCTC8068 | GCA_900475945.1 | Outgroup |
| Pseudomonas_syringae_tagetis_ICMP_4091 | GCF_022557255.1 | Outgroup |
| Pseudomonas_salmasensis_SWRI126 | GCF_014268375.2 | Outgroup |
| Pseudomonas_iranensis_SWRI54 | GCF_014268585.2 | Outgroup |
| Legionella_pneumophila_Philadelphia-1 | GCF_001941585.1 | Outgroup |
| Legionella_micdadei_ATCC_33218 | GCF_000953635.1 | Outgroup |
| Legionella_hackeliae_ATCC_35250 | GCF_000953655.1 | Outgroup |
| Legionella_jordanis_NCTC11533 | GCA_900637635.1 | Outgroup |
| Legionella_fallonii_LLAP-10 | GCF_000953135.1 | Outgroup |
| Legionella_lytica_PCM_2298 | GCF_023921225.1 | Outgroup |
| Legionella_antarctica_TUM19329 | GCF_011764505.1 | Outgroup |
| Legionella_waltersii_NCTC13017 | GCA_900187095.1 | Outgroup |
| Legionella_cardiaca_H63 | GCF_029026145.1 | Outgroup |
| Legionella_spiritensis_NCTC11990 | GCA_900186965.1 | Outgroup |
| Legionella_clemsonensis_CDC-D5610 | GCF_002240035.1 | Outgroup |
| Legionella_lansingensis_NCTC12830 | GCA_900187355.1 | Outgroup |
| Legionella_geestiana_1308 | GCF_004571195.1 | Outgroup |
| Legionella_longbeachae_NSW150 | GCF_000091785.1 | Outgroup |
| Legionella_sainthelensi_LA01-117 | GCF_002848365.2 | Outgroup |
| Legionella_israelensis_L18-01051 | GCF_007361795.1 | Outgroup |
| Candidatus_Legionella_polyplacis_PsAG | GCF_002776555.1 | Outgroup |
| Legionella_anisa_UMCG_3A | GCF_003176875.1 | Outgroup |
| Legionella_oakridgensis_Oak_Ridge-10 | GCF_001467925.1 | Outgroup |
| Legionella_parisiensis_NCTC11983 | GCA_900461585.1 | Outgroup |
